# Learning naturalistic temporal structure in the posterior medial network

**DOI:** 10.1101/196287

**Authors:** Mariam Aly, Janice Chen, Nicholas B. Turk-Browne, Uri Hasson

**Affiliations:** Princeton Neuroscience Institute; Princeton University; Princeton, NJ, 08544; USA; Department of Psychology; Columbia University; New York, NY, 10027; USA; Department of Psychological & Brain Sciences; Johns Hopkins University; Baltimore, MD, 21218 USA; Department of Psychology; Princeton University; Princeton, NJ, 08544; USA; Department of Psychology; Yale University; New Haven, CT, 06520; USA

**Keywords:** temporal regularities, statistical learning, network dynamics, precuneus, hippocampus

## Abstract

The posterior medial network is at the apex of a temporal integration hierarchy in the brain, integrating information over many seconds of viewing intact, but not scrambled, movies. This has been interpreted as an effect of temporal structure. Such structure in movies depends on pre-existing event schemas, but temporal structure can also arise de novo from learning. Here we examined the relative role of schema-consistent temporal structure and arbitrary but consistent temporal structure on the human posterior medial network. We tested whether, with repeated viewing, the network becomes engaged by scrambled movies with temporal structure. Replicating prior studies, posterior medial regions were immediately locked to stimulus structure upon exposure to intact but not scrambled movies. However, for temporally structured scrambled movies, functional coupling within the network increased across stimulus repetitions, rising to the level of intact movies. Thus, temporal structure is a key determinant of network dynamics and function in the posterior medial network.

## Introduction

The brain’s activity is influenced by events across multiple timescales, from seconds to minutes ago (Chaudhuri et al., 2015; Hasson et al., 2015; Murray et al., 2014). Prior studies that have investigated the integration of information over time have often relied on naturalistic stimuli, such as movies or stories (e.g., Hasson et al., 2004, 2008; Honey et al., 2012a,b; Lerner et al., 2011). These stimuli are presented either intact or temporally scrambled (divided into randomly re-ordered segments). Scrambling can be done at different temporal granularities (e.g., at the word, sentence, or paragraph levels), and which level of scrambling disrupts activity in a given brain region indicates its timescale of processing. For example, sensory areas track instantaneous physical parameters (e.g., speech volume) and are unaffected by even the finest scrambling, indicating that they have short processing timescales (Hasson et al., 2008). Conversely, higher-order areas in the posterior medial network (Ranganath and Ritchey, 2012), which overlap with the default mode network (Raichle, 2015), are sensitive to temporal context extending tens of seconds in the past (e.g., a previous sentence can affect the response to an incoming word). They can be disrupted by even coarse scrambling, indicating that they have long processing timescales (Hasson et al., 2015).

Why do higher-order areas respond in a reliable manner to intact, but not scrambled, movies and stories? Both intact and scrambled movies and stories contain events that unfold over time, but only intact stimuli are predictable over long timescales: they share similarities with learned representations of real-life events we repeatedly encounter, which have temporal structure on the order of seconds or minutes. This structure can influence how new information is interpreted, as well as the ability to remember that information and predict upcoming events (Bartlett, 1932; Bower et al., 1979; Gilboa & Marlatte, 2017; van Kesteren et al., 2012). We refer to such events as “schemas” here, but the term “scripts” applies equally, as has been used classically in cognitive psychology to refer to stereotyped or familiar action sequences, like eating in a restaurant (Bower et al., 1979). For example, if food is served to someone at a restaurant, we know that she is having a meal, that she ordered after viewing the menu, and that she will ask for the bill upon finishing. Scrambling the order of events will interfere with our ability to integrate information over time and predict what will happen next, because existing schema no longer apply (e.g., we might see someone order food *after* watching her eat it). Nevertheless, we are capable of learning new event sequences that may initially violate expectations (e.g., paying *before* we eat at a canteen) by extracting temporal regularities across repetitions, a process known as statistical learning (Aslin & Newport, 2012; Schapiro et al., 2017). Here, we explore whether brain regions that accumulate information over long timescales for schema-consistent movies can become engaged by novel event sequences over repeated viewings.

We scanned individuals with fMRI while they watched three 90-s movie clips, six times each, from *The Grand Budapest Hotel* (Figure 1). One clip (“Intact”) was viewed in its original format: a temporally structured clip consistent with pre-existing schemas of how real-life events unfold. Another clip (“Scrambled-Fixed”) comprised short segments that were randomly re-ordered, but their order was identical across all repetitions, creating incoherent but stable temporal structure. A final clip (“Scrambled-Random”) also comprised scrambled segments, but the segments were randomly re-ordered for each repetition, creating incoherent and unstable temporal structure. In contrast to studies in which scrambled movies were presented in the same scrambled order twice (e.g., Hasson et al., 2008; Honey et al., 2012a), the six presentations used here allowed us to search for neural correlates of temporal structure over repeated presentations.

We focused on regions of interest within the posterior medial network (Ranganath & Ritchey, 2012) which overlap with the default mode network, because of their role in processing long-timescale temporal context (Hasson et al., 2015). These regions include the precuneus, the posterior cingulate cortex, and the angular gyrus (Ranganath & Ritchey, 2012). We also included the hippocampus, a region that is part of both the posterior medial and the anterior temporal networks (Ranganath & Ritchey, 2012), because of its role in memory for temporally structured events and its connections to the default mode network (Davachi & DuBrow, 2015; Hasson et al., 2015).

**Figure 1.**
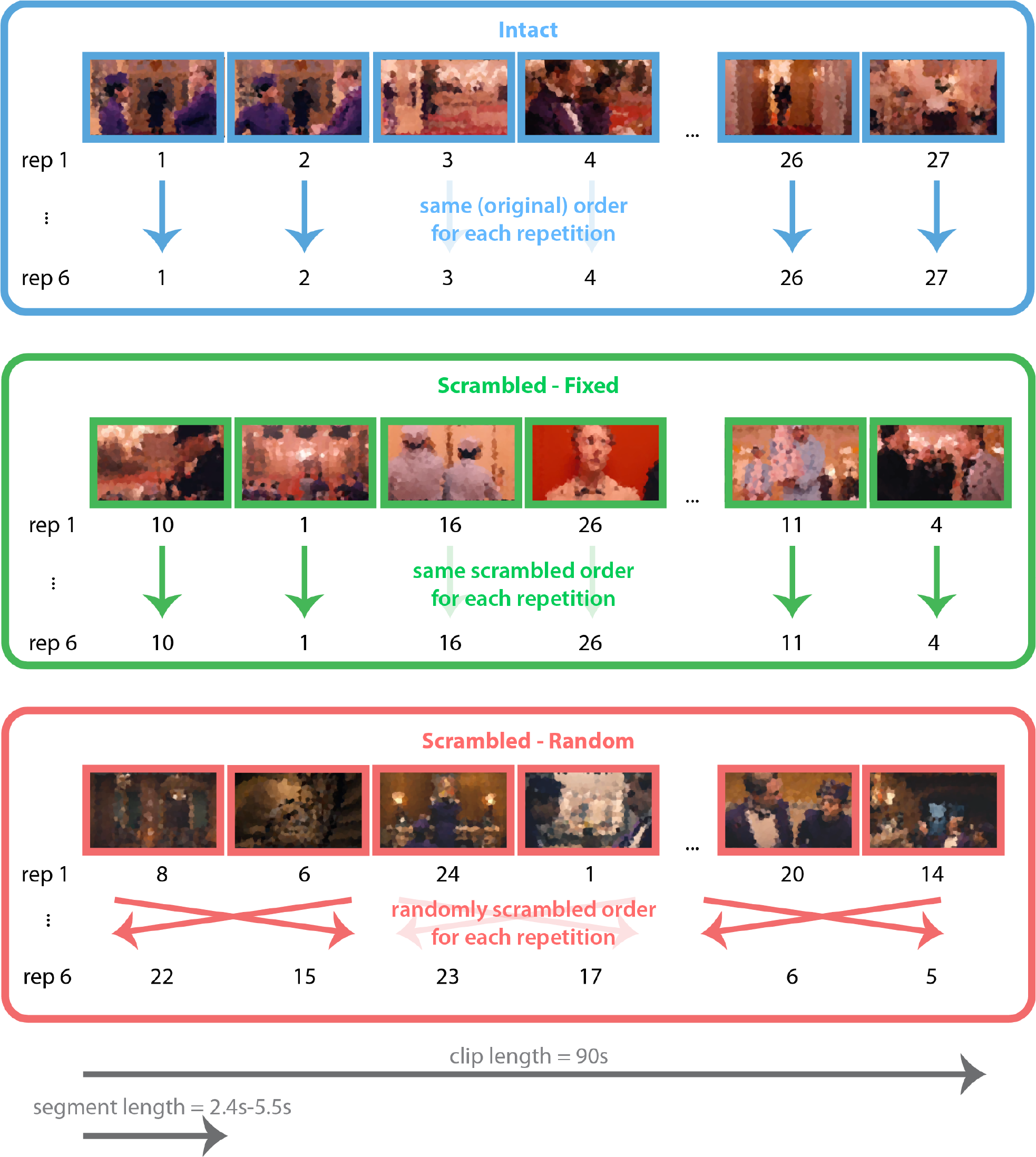
Stimuli and design. Subjects viewed 90-s clips from The Grand Budapest Hotel. Each of three clips was assigned to one of three conditions. Each clip was viewed six times. Numbers below each screenshot refer to a segment within each clip, with segments being continuous chunks 2.4–5.5s long. The Intact clip was viewed in its original format. The segments for the Scrambled-Fixed clip were randomly re-ordered, but viewed in the same scrambled order for all repetitions. The segments for the Scrambled-Random clip were also randomly re-ordered, but the segments were viewed in a different order for each repetition. Movie stills are pixelated in this figure for copyright reasons.

To examine brain activity dynamics for these intact and scrambled movie clips, we conducted several analyses that provide complementary insights (Figure 2; see Hasson et al., 2008; Honey et al., 2012a; Simony et al., 2016). We first examined the consistency of brain activity dynamics within a given brain region in two ways: (1) *intra-subject correlation* — how consistently a given brain region responds in a given individual across different viewings of a move clip (Figure 2A), and (2) *inter-subject correlation* — how consistently a given brain region responds in different individuals watching the same movie clip (Figure 2B).

We also examined the consistency of activity dynamics between different brain regions in two ways: (1) *intra-subject functional correlation* — the coupling in activity between different brain regions in a given individual during a given viewing of a movie clip (Figure 2C), and (2) *inter-subject functional correlation* — the correlation in activity between a brain region in one individual and a different brain region in other individuals watching the same movie clip (Figure 2D).

These analyses, in tandem, can reveal important information about how a stimulus is processed in the brain. High intra-subject correlations, alongside low inter-subject correlations, would indicate that each individual is processing the stimulus in a unique way. Low intra- subject correlations, alongside low inter-subject correlations, may indicate that each individual is processing the stimulus in a unique way that is unstable (i.e., changes) over stimulus repetitions (this is just one possibility, however, as noisy data may produce the same result). Likewise, within-region (inter- and intra-subject correlation) and between-region (inter- and intra-subject functional correlation) analyses offer complementary information because regions can change how they interact with one another whether or not they individually show changes in overall activity (Al-Aidroos et al., 2012; Córdova et al., 2016). For example, increases in intra-subject functional connectivity over stimulus repetitions might indicate that brain regions within an individual start to work in unison as regularities in the stimulus are learned.

**Figure 2.**
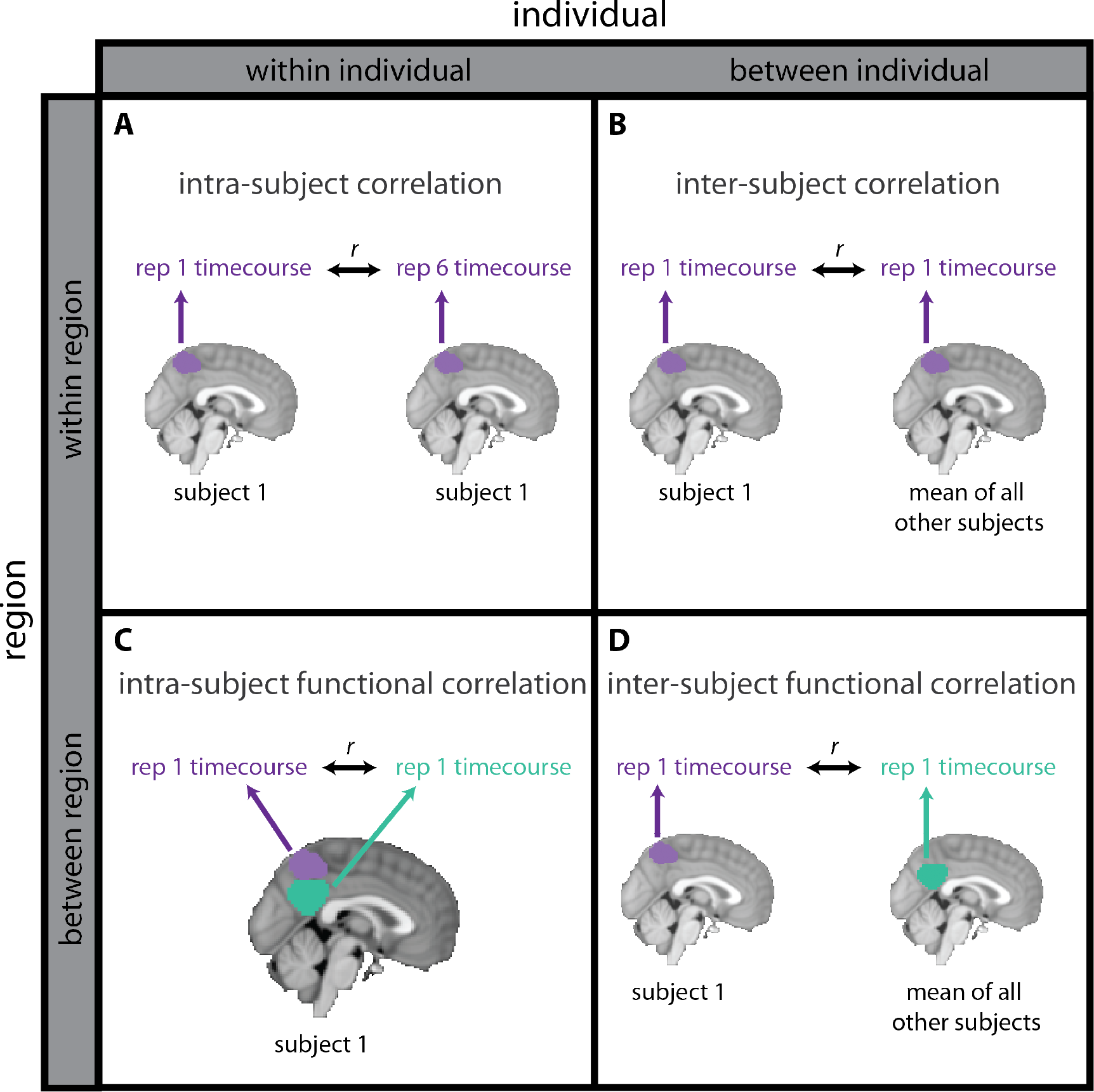
Schematic of fMRI analyses. **(A)** Intra-subject correlation consists of calculating the correlation between a single brain region’s activity timecourses for different repetitions of a given stimulus, within an individual. **(B)** Inter-subject correlation consists of calculating the correlation between a single brain region’s activity timecourse for different individuals watching the same stimulus, for the same repetition of that stimulus. **(C)** Intra-subject functional correlation consists of calculating the correlation between different brain regions’ activity timecourses, within a given stimulus repetition, within an individual. **(D)** Inter-subject functional correlation consists of calculating the correlation between a brain region’s activity timecourse in one individual and a different brain region’s activity timecourse in different individuals, for a given stimulus repetition. For simplicity, only two brains are depicted for the inter-subject analyses, but these analyses involve comparing a given individual to the mean of all other individuals. Inter-subject correlation **(B)** and inter-subject functional correlation **(D)** identify components of brain activity dynamics that are shared between individuals, and thus related to the common stimulus viewed by different individuals. Inter-subject correlation **(B)** identifies such commonalities within a brain region, while inter-subject functional correlation **(D)** identifies commonalities shared between different brain regions. Intra-subject correlation **(A)** and intra-subject functional correlation **(C)** identify components of brain activity dynamics that could either be related to the stimulus being viewed or idiosyncratic to each individual. Thus, comparing inter- and intra-subject analyses offers insights into which components of activity are shared between people (inter-subject analyses) and which may be idiosyncratic to each individual (intra-subject analyses). See Materials and Methods for more details.

In sum, by comprehensively examining temporal dynamics in the posterior medial network over movie clip repetitions, we tested how the network’s responses change with exposure to novel and schematic temporal structure. The critical question concerned whether the posterior medial network can become engaged during viewing of novel temporal structure (in scrambled movies) or only by temporal structure consistent with pre-existing schema (in intact movies) ^1^.

## Materials and Methods

### Subjects

Thirty individuals (12 male) from the Princeton University community participated for monetary compensation (age: *M* = 23.0 years, *SD* = 4.2; education: *M* = 15.3 years, *SD* = 3.2; all right-handed). The sample size was determined *a priori* based on prior fMRI studies using naturalistic stimuli and employing analysis techniques identical to those in the current study (e.g., Andric et al., 2016; Chen et al., 2016, 2017; Hasson et al., 2004, 2008; Honey et al., 2012b; Lerner et al., 2011; Simony et al., 2016). Although the sample size varies across these studies, the average is about 15 subjects. Because we had two counterbalancing conditions (see below), we opted to run 15 subjects in each, for a total sample size of 30.

We only tested individuals who had not previously seen the movie used in this study. Informed consent was obtained from all subjects, and the study was approved by the Institutional Review Board at Princeton University.

### Design and Procedure

Three 90-s clips (referred to as A, B, and C below) were selected from *The Grand Budapest Hotel*, and viewed six times each. We chose six stimulus viewings on the basis of a prior study (Turk-Browne et al., 2010) that found robust statistical learning effects, in the brain and in behavior, after six stimulus repetitions. Each clip included multiple characters, dialogue, music, and movement through several rooms and hallways (for more information about each clip, see Supplementary Methods). Each clip was assigned to one of three conditions (Figure 1). All subjects viewed all three clips. The accompanying audio (dialogue and music) was delivered with MRI-compatible Sensimetrics in-ear headphones. To make the stimuli for each condition, we divided each 90-s clip into short segments, keeping the divisions at natural breaks as much as possible (e.g., an editor’s cut, the end of a spoken sentence). Clip A was divided into 27 segments (2.5–4.4 s). Clip B was divided into 24 segments (2.4–5.5 s). Clip C was divided into 26 segments (2.7–4.6 s).

Although these segments are quite short, comprehensible information is still conveyed within each segment. Each segment included at least one (but usually more than one) meaningful action and/or spoken phrase. Thus, the scrambled movies were not incomprehensible speech and discontinuous movements. Rather, enough information was present in each short segment that subjects were still able to report, after the experiment, what happened in each clip — what was said, what actions were taken, and the general plot (see Behavioral Results) — even for the scrambled movies.

The *Intact* clip was viewed in its original format, as a temporally continuous clip with unfolding events consistent with schemas learned over the course of our lifetime (e.g., what happens when an elevator button is pushed, or what happens when walking into a hotel). This clip was reconstructed in its original order from the shorter segments, so that any audiovisual artifacts introduced by editing would be present for the Intact condition as well as the Scrambled conditions described below. For all subjects, the Intact clip was clip A. For the *Scrambled-Fixed* clip, the segments were randomly re-ordered at first (disrupting consistency with event schemas in semantic memory), but then viewed in the same scrambled order for all six repetitions of the clip (thus, segment transitions were perfectly predictable after the first presentation, providing the basis for learning temporal structure over repetitions). For the *Scrambled-Random* clip, the segments were randomly re-ordered separately for each of the six repetitions of the clip (thus, segment transitions were unpredictable, and segment-level temporal structure was eliminated). There were two counterbalancing conditions: For half of the subjects, the Scrambled-Fixed clip was clip B and the Scrambled-Random clip was clip C, and vice versa for the other half. All subjects in a given counterbalancing condition viewed identical stimuli, i.e., the Scrambled-Fixed clip was scrambled in the same way for all subjects (and for all repetitions), and the Scrambled-Random clip was scrambled in the same way for all subjects for any given repetition (but scrambled differently for different repetitions).

We did not run a fully counterbalanced experiment (i.e., with all possible clip-to-condition assignments). Such a fully counterbalanced experiment would have required an additional 60 subjects (total) across 4 additional counterbalancing orders. The reason we chose to counterbalance as we did is that clip B was quite different from clips A and C: clip B, relative to both clips A and C, had fewer characters, softer and slower music, and a slower pace of action (we tried to match the clips as much as possible, but were not able to in this regard). Thus, we counterbalanced the assignment of clips B and C to the scrambled conditions because we did not want statistically significant effects in the Scrambled-Fixed, but not the Scrambled-Random, condition to be a result of the movie clips assigned to those conditions.

To summarize the design: The Intact and Scrambled-Fixed clips were both repeated identically for all six presentations of each clip. The former was in its original, continuous format; thus, temporal expectations existed on the very first exposure because of consistency with real-world event schemas. The latter became temporally predictable over repetitions because of deterministic transitions between re-ordered movie segments. The Scrambled-Fixed and Scrambled-Random clips both contained random events inconsistent with real-world predictions, but the latter was viewed in a different scrambled order every time. Although the Scrambled-Random clip contained temporal structure on short timescales (e.g., within the ~3s duration of each segment), longer timescale structure across segments was present only in the Intact and Scrambled-Fixed clips.

The experiment was divided into three runs, each ~10 minutes long. Each run began with 4.5 s of blank time and then an additional 6-s countdown screen to alert subjects to the upcoming start of the movie clips. The rest of each run comprised six movie clips in total (clips A, B, and C viewed twice each), with 3 s of fixation and then 6 s of countdown between each clip. Across runs, A, B, and C were each viewed in each of the six serial positions in a run (e.g., A was viewed in positions one and five in run one, positions two and four in run two, and positions three and six in run three). The same clip was never viewed back-to-back within a run. The within-run lag between repetitions of a clip was matched across the clips: in one run, there was a lag of one between repetitions, in another there was a lag of two, and in another there was a lag of three between repetitions of the same clip. All subjects viewed A, B, and C in the same order, but the assignment of clip to condition differed across subjects.

Subjects were instructed to view the clips, but there was no other task during the scan. It was emphasized that attention should be paid at all times. To verify this, we surprised subjects with a post-scan free recall test, which required them to type everything they could remember about each of the clips. Subjects were generally able to recall the main events and characters in each clip and (varying amounts of) perceptual details (see Behavioral Results).

### fMRI Methods

#### Acquisition

Data were collected on a 3T Siemens Prisma scanner with a 64-channel head/neck coil. Functional images were obtained with a multiband EPI sequence (TR = 1.5 s, TE = 39 ms, flip angle = 50°. acceleration factor = 4, shift = 3, voxel size = 2.0-mm iso), with 60 oblique axial slices acquired in an interleaved order. Whole-brain high-resolution (1.0-mm iso) T1-weighted structural mages were acquired with an MPRAGE sequence. Field maps were collected for registration, consisting of 40 oblique axial slices (3-mm iso).

#### Software

Preprocessing, registration, and permutation tests for whole-brain analyses were conducted using FEAT, FLIRT, and command-line functions in FSL (http://fsl.fmrib.ox.ac.uk/fsl/). All other analyses were performed with custom Matlab scripts.

#### Preprocessing

The first three volumes of each run were discarded to allow for T1 equilibration. Preprocessing steps included brain extraction, motion correction, high-pass filtering (max period = 140 s), spatial smoothing (3-mm FWHM Gaussian kernel), and registration to standard MNI space (via an intermediate registration to the subject’s anatomical image with boundary-based registration).

The fields maps were preprocessed using a custom script, in accordance with the FUGUE user guide from FSL. First, the two field map magnitude images were averaged together and skull stripped. The field map phase image was converted to rad/s and smoothed with a 2-mm Gaussian kernel. The resulting phase and magnitude images were included in the preprocessing step of FEAT analyses to unwarp the functional images and aid registration to anatomical space. Following registration, the distortion-corrected functional images were compared to the originals to ensure that unwarping was effective. In all cases, using the field maps reduced distortion in anterior temporal and frontal regions.

After preprocessing, the filtered 4D functional images for each run were divided into volumes that corresponded to each of the six clips presented within that run. Four volumes were then cropped from the beginning of each clip to adjust for hemodynamic lag. Finally, the timecourse for each clip was z-scored within each subject and region of interest (ROI).

#### ROI definition

We focused our analyses primarily on the precuneus, as a key node in the posterior medial (and default mode) network (Hasson et al., 2015; Ranganath and Ritchey, 2012; Utevsky et al., 2014), and a region that is consistently and robustly recruited by long-timescale regularities (Baldassano et al., 2017; Chen et al., 2016, 2017; Hasson et al., 2010, 2015; Honey et al., 2012b; Lerner et al., 2011). In addition to examining activity within the precuneus, we investigated its interactions with three other ROIs in the posterior medial network: the hippocampus, angular gyrus, and posterior cingulate cortex (Ranganath & Ritchey, 2012); we could not include medial prefrontal cortex because of excessive signal dropout (we oriented our slices parallel to the long axis of the hippocampus to maximize coverage of the hippocampus and parietal cortex, our main regions of interest; unfortunately, this slice orientation increases signal dropout in medial prefrontal cortex, which is susceptible to dropout because of its proximity to the nasal sinuses). For completeness, we also examined all pairwise interactions between the four posterior medial ROIs. These ROIs were defined bilaterally with the Harvard-Oxford Structural Atlas. The precuneus and posterior cingulate ROIs were edited to not contain overlapping voxels, with the precuneus ROI comprising dorsomedial parietal cortex and the posterior cingulate ROI comprising ventromedial parietal cortex.

We also examined a control ROI in early visual cortex, chosen for its relatively short temporal integration window in prior studies (e.g., Chen et al., 2016; Hasson et al., 2008). The ROI was defined functionally from a separate group of 25 subjects watching two minutes of an audiovisual movie (Chen et al., 2016), including voxels bilaterally along the collateral sulcus with the highest inter-subject correlation (*r*s > 0.45).

The timecourse of activity was averaged across all voxels in an ROI, and these mean ROI timecourses were submitted to further analysis. We conducted within-region and between-region analyses, within-individual (intra-subject) and between individuals (inter-subject), for complementary insights into how the brain changes with repeated exposure to stimuli that vary in temporal structure. These analyses, along with their logic, are described in more detail below.

#### Within-region correlations

We examined the consistency of BOLD responses to the movie clips in two ways, both of which have been used in prior studies of temporal integration with naturalistic stimuli (e.g., Hasson et al., 2005, 2008; Honey et al., 2012a,b; Lerner et al., 2011): (1) intra-subject correlation over clip repetitions (Figure 2A), and (2) inter-subject correlation for the same clip repetition (Figure 2B). As noted above, for these analyses we focused on the precuneus; subsequent analyses incorporated additional regions and whole-brain analyses.

To assess the consistency of precuneus activity to a given clip within an individual (i.e., intra-subject correlation), the timecourse of activity for repetition 1 of each clip was compared to that of repetition 6 of the same clip using Pearson correlation^2^. For the Intact and Scrambled-Fixed clips, the order of segments within each clip was the same across repetitions, thus brain activity was being compared for objectively identical stimuli in repetitions 1 and 6. For the Scrambled-Random clip, segment order was different across all repetitions, thus brain activity was being compared for *different* stimuli in repetitions 1 and 6. Thus, this condition serves as a baseline because correlations should approach zero even in early sensory areas.

Planned comparisons between conditions were conducted at the group level with random-effects paired *t*-tests, after Fisher transformation of the correlation coefficients to ensure normality. Two-tailed *p*-values and 95% confidence intervals are reported for this and all other analyses, as well as the correlation values as a measure of effect size.

We also performed this analysis for the whole brain. We calculated, for each voxel in the brain for each individual, the correlation between the timecourse of activity for repetition 1 of each clip and repetition 6 for each clip. These whole-brain intra-subject correlation maps were analyzed at the group level using random-effects non-parametric tests (randomise in FSL) and corrected for family-wise error (FWE) across all voxels. The resulting maps were thresholded at *p* < 0.05, FWE corrected (Supplementary Figure S1).

To assess the consistency of precuneus activity to a given clip across individuals (i.e., inter-subject correlation), we correlated each subject’s timecourse of activity for repetition 1 of each clip with the mean of all *other* subjects’ timecourses for that same repetition and clip. Because subjects viewed the same stimulus but may have had different thoughts and emotions triggered by it, correlating timecourses across individuals highlights the stimulus-locked brain activity that is shared across people irrespective of these individual differences. In contrast, the intra-subject correlation analysis described above can identify idiosyncratic responses that are particular to a person and/or asynchronous across people (Hasson et al., 2004, 2009).

For this analysis, we focused on the first time each clip was viewed because this allowed a measure of the consistency of precuneus dynamics between subjects upon initial exposure, without any influence of memory for previous times the clip was viewed. This additionally allows us to compare our results to published studies with only one presentation of a movie clip (e.g., Chen et al., 2016; Hasson et al., 2004; Lerner et al., 2011). Nevertheless, for completeness, we also examined the inter-subject correlation for the last repetition of each clip (Supplementary Figure S2).

There were two clip-to-condition counterbalancing subgroups (i.e., clip B was the Scrambled-Fixed clip for half of subjects and Scrambled-Random for the other half), so inter-subject correlation was calculated within each subgroup and then pooled. In this way, we obtained the correlation between brain activity of different people viewing identical stimuli (this includes identical segment ordering for the scrambled clips). Specifically, the brain activity of each of the subjects numbered 1-15 was correlated against the average of the other 14 subjects numbered 1-15 (these subjects watched identical clips). Likewise, the brain activity of each of the subjects numbered 16-30 was correlated against the average of the other 14 subjects numbered 16-30 (these subjects watched identical clips). Thus, each subject had a correlation value for his/her brain activity vs. the mean of all other subjects in his/her counterbalancing group. All 30 correlations were then used for group analyses. We conducted inter-subject analyses for the Intact clip within the two subgroups of 15 subjects in the same manner (even though all subjects saw the same clip). Thus, this analysis was consistent and equivalently powered across conditions.

The resulting inter-subject correlation data are not independent, however: For *n* subjects in the same counterbalancing group, any pair of subjects will share *n* – 2 timecourses in the calculation of the mean “others” timecourse. For example, the “others” timecourse for subject 1 contains data from subjects 2-15, and the “others” timecourse for subject 2 contains data from subjects 1, 3-15. Thus, data for subjects 3-15 were used to calculate the inter-subject correlation for both subject 1 and subject 2. The inter-subject correlation values for different subjects are therefore not independent. Because of this, we conducted non-parametric permutation tests to compare the mean inter-subject correlation for different conditions. For each subject, we took the difference between the two conditions of interest (after Fisher-transforming the correlations) and then flipped the sign of the difference (effectively swapping condition assignments) for some random subset of subjects before obtaining the group mean. This was repeated 1000 times, and the distribution of mean differences for the randomly flipped data was compared to the observed mean difference between conditions to obtain *p*-values. Two-tailed *p*-values were obtained by finding the smaller tail of the null distribution vs. the observed mean and multiplying that value by two.

In addition to this approach, we also calculated confidence intervals (CIs) on the mean difference between conditions using a bootstrap approach. For each subject, we took the difference between the two conditions of interest (after Fisher-transforming the correlations), randomly sampled subjects with replacement, and calculated the mean from this sample. This was repeated 1000 times, and the points below and above which 2.5% of the samples lay (i.e., the bounds that contained 95% of the data) were the bootstrap confidence intervals. Both of these approaches (i.e., the permutation test and the bootstrapped CIs) yielded comparable results to paired-samples *t-*tests.

#### Between-region correlations

Intra-subject functional correlation (Figure 2C) between ROIs was examined across all repetitions of each clip to see how neural coupling changes with stimulus repetitions. For each repetition and clip, the timecourse of BOLD activity in the precuneus was correlated with that of the other ROIs, for the same repetition and clip. After Fisher transformation, four types of analyses were conducted to assess changes in intra-subject functional correlation: (1) a 2 × 3 repeated-measures analysis of variance (ANOVA) with repetition (1 or 6) and condition (Intact, Scrambled-Fixed, Scrambled-Random) as factors; (2) planned comparisons of repetitions 1 vs. 6 within condition using paired *t*-tests; (3) a 6 × 3 repeated-measures ANOVA with repetition (1-6) and condition (Intact, Scrambled-Fixed, Scrambled-Random) as factors; and (4) a test of whether the slope across repetitions 1-6 within condition was different from zero, using one-sample *t-*tests.

Together, these four analyses offered a complete picture of changes in functional correlation as movies are repeated: Analyses 1 and 2 allowed us to examine intra-subject functional correlation at the beginning vs. end of exposure (repetitions 1 vs. 6), whereas analyses 3 and 4 allowed us to examine the nature of change over the course of all six repetitions. The ANOVAs (analyses 1 & 3) allowed us to examine whether differences between conditions and repetitions existed, whereas the more-targeted within-condition analyses (analyses 2 & 4) allowed us to examine changes in brain activity in each condition separately.

For completeness, we also examined intra-subject functional correlation between all pairs of posterior medial ROIs. To visualize changes in functional coupling, we made network graphs for the first vs. last repetitions using a Matlab toolbox *circularGraph*, where each posterior medial ROI was a node, and the edges represented the strength of coupling between all pairs of ROIs.

To assess the selectivity of changes in intra-subject functional correlation within the posterior medial ROIs, we conducted two additional analyses. First, we examined intra-subject functional correlation between the precuneus and a control ROI in early visual cortex. Second, we conducted whole-brain, voxelwise intra-subject functional correlation analyses using the precuneus as the seed ROI. For each subject, clip, and repetition, the timecourse of BOLD activity in the precuneus was correlated with the timecourse of every other voxel in the brain. A contrast map of repetition 6 minus repetition 1 was generated for each subject and condition. Within each condition, these contrast maps were analyzed at the group level using random-effects non-parametric tests (randomise in FSL), and corrected for family-wise error (FWE) across all voxels. The resulting maps were thresholded at *p* < 0.05 corrected.

We also conducted functional correlation analyses between subjects (Figure 2D). Inter-subject functional correlation (Simony et al., 2016) is analogous to the inter-subject correlation analyses described above, but the timecourse of activity for a given ROI and subject is correlated to the mean of all other subjects’ timecourses for a different ROI (within repetition and condition). For example, to calculate inter-subject functional correlation between the hippocampus and precuneus, we first obtained the correlation between the hippocampal timecourse for a particular individual and the mean precuneus timecourse across all other individuals. Then, we calculated the correlation between the precuneus timecourse for the same individual and the mean hippocampal timecourse across all other individuals. The average of these calculations is taken as the measure of hippocampus-precuneus inter-subject functional correlation for that subject. The analysis is repeated for all subjects, and the resulting data are submitted to group-level analysis. As with the inter-subject correlation analysis, we calculated inter-subject functional correlation within each subgroup of 15 subjects that viewed identical stimuli, and then pooled across subjects. The resulting data were analyzed using permutation tests and bootstrap methods, as above.

## Results

### Behavioral Data

Free recall data were scored by counting the number of details reported by each subject for each movie clip. Details included (but were not limited to) characters, dialogue, actions, and perceptual details such as the appearance of characters and the scenes through which they moved. Reported details were checked against the movie clips for veracity.

Subjects recalled an average of 25.75 details for the Intact clip (SD = 11.24), 21.45 details for the Scrambled-Fixed clip (SD = 12.62), and 19.93 details for the Scrambled-Random clip (SD = 8.82). Memory for the Intact clip was better than memory for the Scrambled-Fixed clip (*t*_29_ = 2.17, *p* = .04, 95% CI: 0.24–8.36) and the Scrambled-Random clip (*t*_29_ = 3.36, *p* = .002, 95% CI: 2.28–9.36). Recall scores for the Scrambled-Fixed and Scrambled-Random clips were not significantly different (*t*_29_ = 0.66, *p* = .51, 95% CI:-3.17–6.20).

In their recalls for the scrambled movie clips, subjects re-ordered the clip segments into a coherent narrative. Because of this, and because we did not have a separate temporal order memory test for the movie clip segments, we cannot directly speak to learning of temporal structure as it may be evidenced in behavior. Our focus in this paper, however, is learning of temporal structure as it may be evidenced in the brain.

### Within-Region Correlations

Prior studies have found that temporal dynamics within posterior medial regions, such as the precuneus, are more consistent for intact vs. scrambled movies and stories. Our first aim was to replicate these findings — specifically, to show that temporal fluctuations in the precuneus are more highly correlated for intact vs. scrambled stimuli when measured: (1) within an individual across repetitions of these clips, i.e., intra-subject correlation (Figure 2A; e.g., Hasson et al., 2004, 2008; although only two repetitions were used in that study), and (2) between individuals upon initial exposure to these clips, i.e., inter-subject correlation (Figure 2B; e.g., Honey et al., 2012b; Lerner et al., 2011).

#### Consistency Within Each Individual

For the intra-subject correlation analysis, it is important to note that the Scrambled-Random clip was an objectively different stimulus on the first and last repetitions (i.e., the segments within the clip were presented in a different order), but the Intact and Scrambled-Fixed stimuli were objectively identical on the first and last repetition. Thus, the intra-subject correlation analysis for the Scrambled-Random condition is expected to yield low correlations due to changes in the stimulus across runs. The intra-subject correlation for the Scrambled-Random clip is therefore presented here only for completeness, as it is not expected to be significantly different from zero.

**Figure 3.**
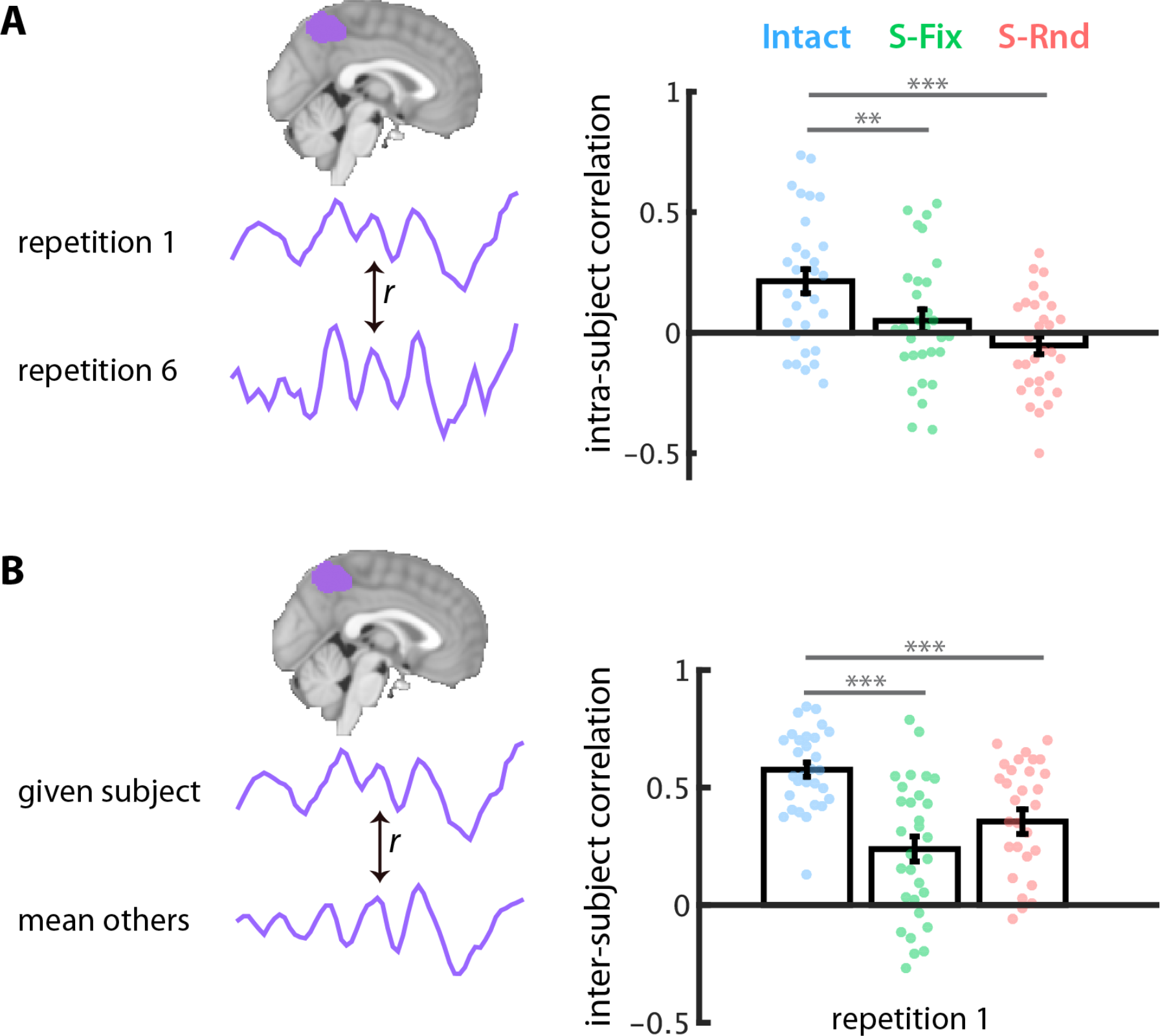
Temporal dynamics in the precuneus are more consistent for intact vs. scrambled movies. **(A)** The timecourses of BOLD activity in the precuneus for the first and last repetitions of each clip were extracted and correlated within subject. The mean intra-subject correlation was higher for the Intact clip compared to the Scrambled-Fixed and Scrambled-Random clips, which did not themselves differ. **(B)** Reliability was also assessed by correlating, for the first repetition, each subject’s timecourse of BOLD activity with the mean of all other subjects. The mean inter-subject correlation was higher for the Intact clip compared to the Scrambled-Fixed and Scrambled-Random clips, which did not themselves differ. Dots indicate individual subjects. Error bars are ± 1 SEM. ** p < .01, *** p < .001.

As shown in Figure 3A, and consistent with prior studies, there was higher intra-subject correlation between precuneus activity for the first and last repetitions of the Intact clip compared to the correlation between the first and last repetitions of the Scrambled-Fixed clip (*r*_intact_ = 0.21, *r*_S-Fix_ = 0.05; *t*_29_ = 2.78, *p* = .009, 95% CI: 0.05–0.32). The correlation for the Intact clip was also higher than that for the Scrambled-Random clip (*r*_intact_ = 0.21, *r*_S-Rnd_ = −0.05; *t*_29_ = 4.26, *p* = .0002, 95% CI: 0.15–0.44), although, as noted above, the correlation for the Scrambled-Random clip was expected to be near zero because the stimulus is different on the first vs. last repetition. Response reliability for the Intact clip was also significantly greater than zero (*t*_29_ = 4.13, *p* = .0003, 95% CI: 0.12–0.36).

Next, we compared the reliability of intra-subject precuneus dynamics for the Scrambled-Fixed and Scrambled-Random clips. Neither of these clips were consistent with pre-existing event schemas, but the Scrambled-Fixed clip contains predictable segment sequences over repetitions (and thus, temporal structure on the order of tens of seconds). However, neural learning of temporal structure in the Scrambled-Fixed clip would not be expected to increase the across-repetition reliability of a region: in fact, the region should be *less* similar to itself over repetitions if learning changes how it processes information. Intra-subject correlations over repetitions should also be low for the Scrambled-Random clip, but for a different reason: because the clip is viewed in a different order every time, there is a different stimulus for repetitions 1 and 6. Indeed, the intra-subject correlation between precuneus activity in repetitions 1 and 6 was not different from zero in either condition (Scrambled-Fixed: *r* = 0.05; *t*_29_ = 1.10, *p* = .28, 95% CI: −0.05–0.16; Scrambled-Random: *r* = −0.05 *t*_29_ = 1.43, *p* = .16, 95% CI: −0.13–0.02) and the conditions were not different from one another (*t*_29_ = 1.64, *p* = .11, 95% CI: −0.03–0.25).

#### Consistency Between Individuals

A similar pattern of results was obtained when examining the reliability of precuneus activity between subjects on the first repetition of each clip (Figure 3B; for the same analysis for the last repetition of each clip, see Supplementary Figure S2). In this case, we examined inter-subject correlation between individuals watching identical stimuli, i.e., the scrambled movies were watched in the same scrambled order for these different subjects. The inter-subject correlation for the Intact clip was higher than that for the Scrambled-Fixed clip (*r*_intact_ = 0.58, *r*_S-Fix_ = 0.24; 95% bootstrapped CI: 0.28–0.57; *p* = 0 from permutation test) and the Scrambled-Random clip (*r*_intact_ = 0.58, *r*_S-Rnd_ = 0.36; 95% bootstrapped CI: 0.19–0.40; *p* = 0 from permutation test). The inter-subject correlation for the Scrambled-Fixed clip was marginally lower than that for the Scrambled-Random clip (95% bootstrapped CI: −0.27–0.01; *p* = .09 from permutation test). However, because the stimuli in these conditions did not differ in the first repetition (i.e., before the scrambled order had been repeated), this difference is not meaningful. Finally, inter-subject correlation in the precuneus was reliably above zero for all three conditions, as expected because all subjects being compared viewed identical stimuli in all conditions (all *p*s = 0 from permutation tests).

Together, these within-region analyses replicate and extend prior work showing greater response reliability in posterior medial regions for intact vs. scrambled movies and stories (e.g., Hasson et al., 2008; Honey et al., 2012a,b; Lerner et al., 2011).

### Between-Region Correlations

#### Within-Subject ROI Analyses

Low inter-subject correlation (Figure 3B), as observed for the Scrambled-Fixed and Scrambled-Random clips, indicates that temporal dynamics in the precuneus are not locked to the external stimulus that is viewed by all subjects. However, intra-subject dynamics were also unstable (Figure 3A). Changing temporal dynamics over clip repetitions could indicate learning in the brain, but they can also occur for other reasons (e.g., fatigue, noisy data). Thus, to further search for learning effects in the brain, we examined network functional correlations during each clip repetition within each individual (c.f. Andric et al., 2016). Analyses of intra-subject functional correlation (Figure 2C) can be useful for understanding how brain regions respond to naturalistic stimuli. For example, functional coupling between the hippocampus and medial prefrontal cortex is modulated by prior exposure to schema-relevant information while watching an intact movie (van Kesteren et al., 2010). We hypothesized that neural learning of temporal structure in the Scrambled-Fixed clip would manifest as enhanced intra-subject functional coupling, over clip repetitions, between the precuneus and other regions in the posterior medial network: hippocampus, angular gyrus, and posterior cingulate cortex (Figure 4).

**Figure 4.**
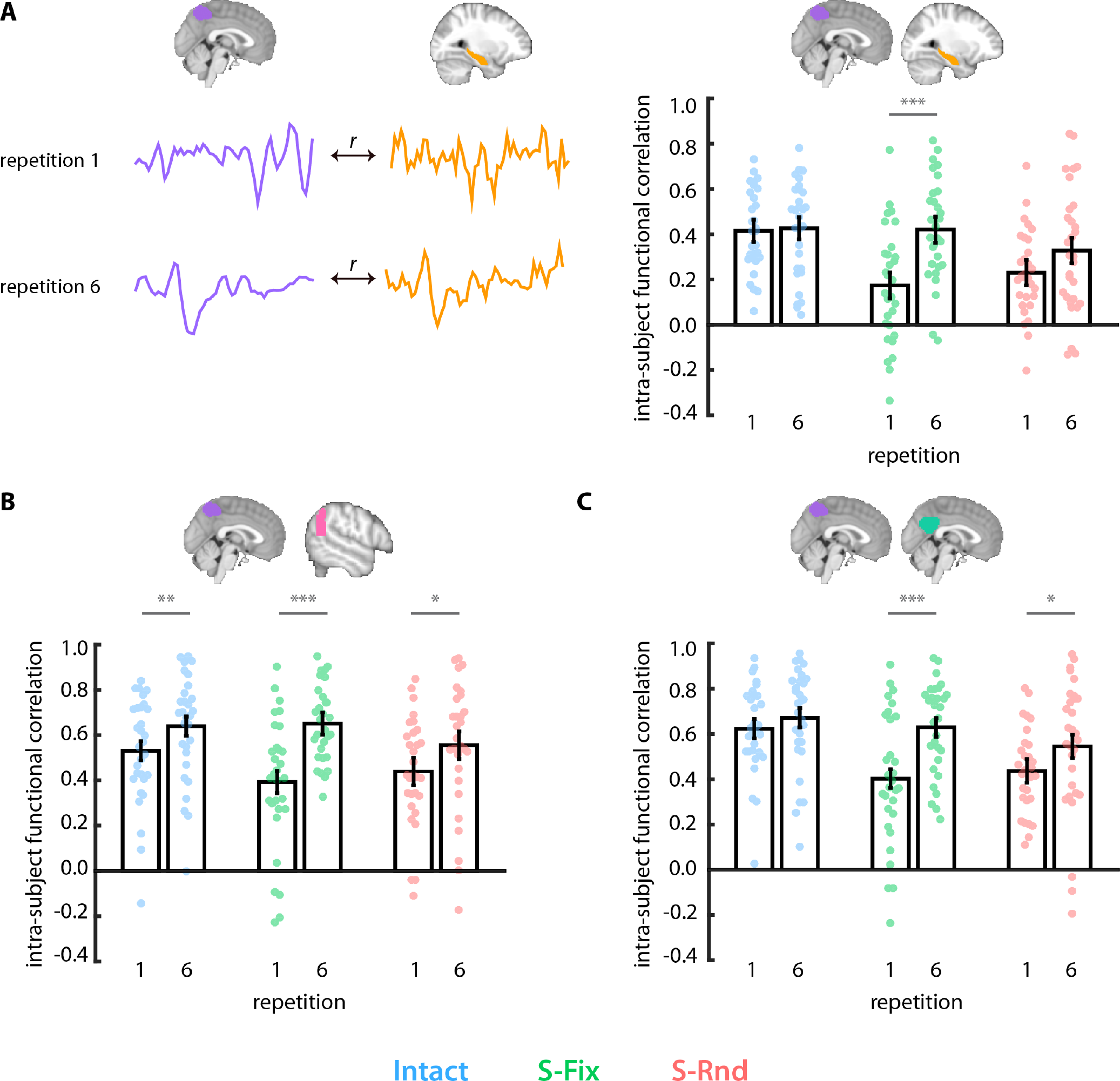
Temporal structure enhances intra-subject functional correlation within the posterior medial network. **(A)** The timecourse of BOLD activity in the precuneus was correlated within-subject with that of the hippocampus for the first and last repetition of each clip. Functional correlations did not change over repetitions of the Intact or Scrambled-Random clips, but more than doubled over repetitions of the Scrambled-Fixed clip, rising to the level of the Intact clip. A similar pattern of results was obtained for **(B)** the precuneus and angular gyrus and **(C)** the precuneus and posterior cingulate. Dots indicate individual subjects. Error bars are ± 1 SEM of the within-subject difference between repetitions. * p < .05, ** p < .01, *** p < .001.

We expected that changes in network intra-subject functional correlations over repetitions would differ between the three conditions. To verify this, we conducted 2 (repetition: 1 or 6) by 3 (condition: Intact, Scrambled-Fixed, Scrambled-Random) repeated-measures ANOVAs, looking specifically for an interaction. Such an interaction was found for coupling between hippocampus and precuneus (*F*_2, 58_ = 6.12, *p* = .004). Follow-up planned comparisons revealed a reliable increase in intra-subject functional correlation from repetition 1 to 6 for the Scrambled-Fixed clip (*r*_Rep1_ = 0.17, *r*_Rep6_ = 0.42, *t*_29_ = 4.38, *p* = .0001, 95% CI: 0.16–0.43) but not the Intact clip (*r*_Rep1_ = 0.42, *r*_Rep6_ = 0.43, *t*_29_ = 0.33, *p* = .74, 95% CI: −0.11– 0.15) or the Scrambled-Random clip (*r*_Rep1_ = 0.23, *r*_Rep6_ = 0.32, *t*_29_ = 1.98, *p* = .06, 95% CI: −0.005–0.28). Indeed, while intra-subject functional correlation was initially higher for the Intact clip than the Scrambled-Fixed clip in repetition 1 (*r*_Intact_ = 0.42, *r*_S-Fix_ = 0.17, *t*_29_ = 5.08, *p* = .00002, 95% CI: 0.16–0.38), this difference disappeared by repetition 6 (*r*_Intact_ = 0.43, *r*_S-Fix_ = 0.42, *t*_29_ = 0.0001, *p* = .9999, 95% CI: −0.13– 0.13).

To obtain additional evidence that increased intra-subject functional correlation from the first to last repetition of the Scrambled-Fixed clip reflected learning in the brain, we examined these functional correlations across all six repetitions. If these changes are in fact indicative of learning, enhancements in intra-subject functional correlation should be monotonic, increasing gradually with exposure to the temporal structure (Supplementary Figure S3). We conducted a 6 (repetition: 1-6) by 3 (condition: Intact, Scrambled-Fixed, Scrambled-Random) repeated-measures ANOVA, which again revealed a significant interaction for precuneus-hippocampus intra-subject functional correlation (*F*_10, 290_ = 2.82, *p* = .002). This interaction was driven by a significant monotonic increase in intra-subject functional correlation over repetitions for the Scrambled-Fixed clip, manifested as a slope greater than zero (β = 0.05??*t*_29_ = 4.06, *p* = .0003, 95% CI: 0.02–0.07). The slope was not different from zero for the Intact clip (β = 0.004, *t*_29_ = 0.43, *p* = .67, 95% CI: −0.02–0.02) and was numerically smaller for the Scrambled-Random clip (β = 0.03, *t*_29_ = 2.17, *p* = .04, 95% CI: 0.002–0.06).

Neither ANOVA revealed a significant repetition x condition interaction for precuneus-angular gyrus intra-subject functional correlation (repetitions 1 vs. 6: *F*_2, 58_ = 1.87, *p* = .16; repetitions 1 through 6: *F*_10,_ 290 = 0.96, *p* = .47) or precuneus-posterior cingulate cortex intra-subject functional correlation (repetitions 1 vs. 6: *F*_2, 58_ = 2.08, *p* = .13; repetitions 1 through 6: *F*_10, 290_ = 1.78, *p* = .06). Nevertheless, within the Scrambled-Fixed condition, intra-subject functional correlation increased from repetitions 1 vs. 6 (precuneus-angular gyrus: *r*_Rep1_ = 0.39, *r*_Rep6_ = 0.65, *t*_29_ = 5.20, *p* = .00001, 95% CI: 0.23–0.54; precuneus-posterior cingulate cortex: *r*_Rep1_ = 0.40, *r*_Rep6_ = 0.63, *t*_29_ = 5.24, *p* = .00001, 95% CI: 0.20–0.46) and the slope across all six repetitions was greater than zero (precuneus-angular gyrus: β = 0.07, *t*_29_ = 4.68, *p* = .00006, 95% CI: 0.04–0.09; precuneus-posterior cingulate cortex: β = 0.06, *t*_29_ = 4.76, *p* = .00005, 95% CI: 0.03– 0.08). The interactions failed to reach significance because of (numerically smaller) increases in precuneus-angular gyrus intra-subject functional correlation for both the Intact clip (repetitions 1 vs. 6: *r*_Rep1_ = 0.53, *r*_Rep6_ = 0.64, *t*_29_ = 3.23, *p* = .003, 95% CI: 0.08–0.37; slope across repetitions: β = 0.04, *t*_29_ = 3.43, *p* = .002, 95% CI: 0.02–0.06) and the Scrambled-Random clip (repetitions 1 vs. 6: *r*_Rep1_ = 0.44, *r*_Rep6_ = 0.56, *t*_29_ = 2.42, *p* = .02, 95% CI: 0.03–0.40; slope across repetitions: β = 0.04, *t*_29_ = 2.42, *p* = .02, 95% CI: 0.007– 0.08), and in precuneus-posterior cingulate cortex intra-subject functional correlation for the Scrambled-Random clip (repetitions 1 vs. 6: *r*_Rep1_ = 0.44, *r*_Rep6_ = 0.55, *t*_29_ = 2.62, *p* = .01, 95% CI: 0.05–0.41; slope across repetitions: β = 0.04, *t*_29_ = 2.48, *p* = .02, 95% CI: 0.007–0.08).

As with precuneus-hippocampus intra-subject functional correlation, both precuneus-angular gyrus intra-subject functional correlation and precuneus-posterior cingulate intra-subject functional correlation were higher for the Intact clip compared to the Scrambled-Fixed clip on repetition 1 (precuneus-angular gyrus: *r*_Intact_ = 0.53, *r*_S-Fix_ = 0.39, *t*_29_ = 2.20, *p* = .04, 95% CI: 0.01–0.34; precuneus-posterior cingulate cortex: *r*_Intact_ = 0.62, *r*_S-Fix_ = 0.40, *t*_29_ = 3.83, *p* = .0006, 95% CI: 0.14–0.48), but this difference disappeared by repetition 6 (precuneus-angular gyrus: *r*_Intact_ = 0.64, *r*_S-Fix_ = 0.65, *t*_29_ = 0.27, *p* = .79, 95% CI: −0.14–0.18; precuneus-posterior cingulate cortex: *r*_Intact_ = 0.67, *r*_S-Fix_ = 0.63, *t*_29_ = 1.49, *p* = .15, 95% CI: −0.04–0.26).

We visualized these effects (and others not reported above) by plotting graphs of all possible pairwise relationships between posterior medial ROIs (Figure 5). There was overall increased functional coupling in the network from repetitions 1 to 6 of the Scrambled-Fixed clip, but less consistent enhancement for the Intact and Scrambled-Random clips.

**Figure 5.**
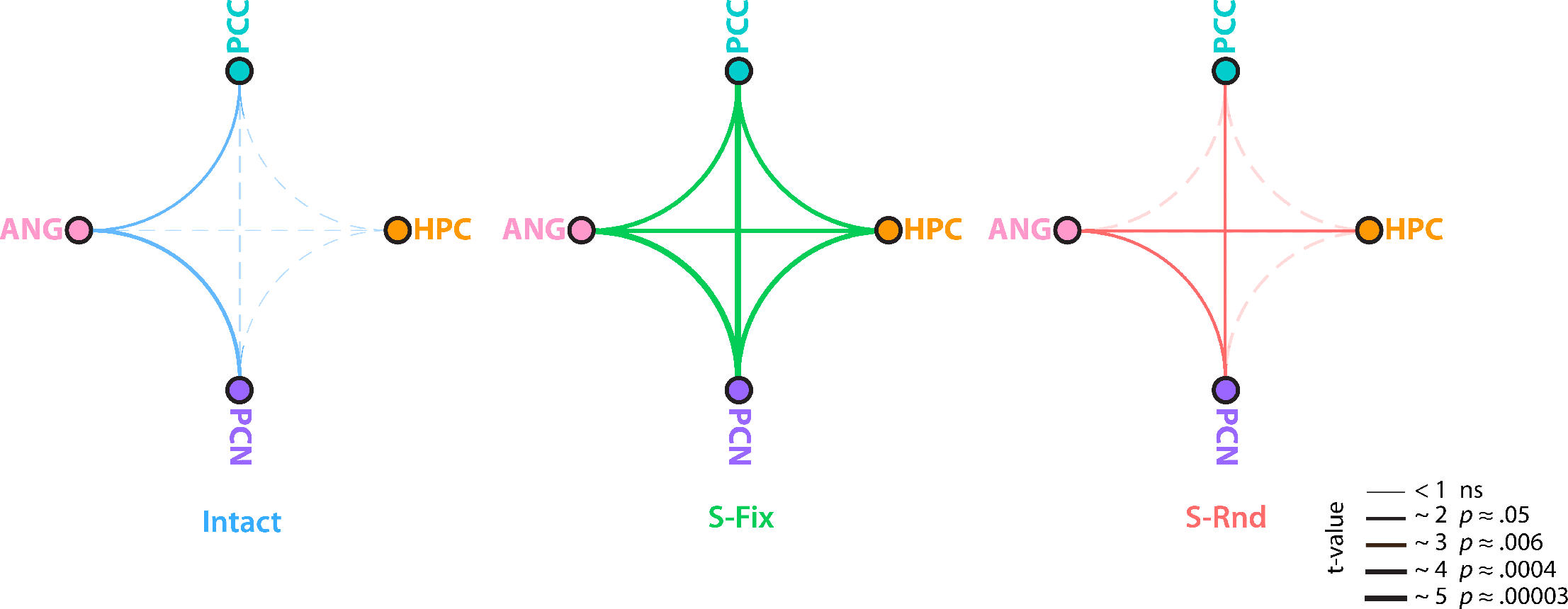
Changes in posterior medial network functional correlations from first to last repetition. Graph visualization of intra-subject functional correlation between all pairs of posterior medial ROIs, including the results from Figure 4, as well as pairs that did not include precuneus. The edges depict the t value comparing intra-subject functional correlation in repetitions 1 vs. 6, with thicker lines indicating greater enhancement over time. Solid lines are statistically significant changes, whereas dashed lines are not statistically significant. The most consistent enhancement in intra-subject functional correlation was for the Scrambled-Fixed clip (all six edges were statistically significant), compared to the Intact clip (two statistically significant edges) and the Scrambled-Random clip (three statistically significant edges). PCC = posterior cingulate cortex, HPC = hippocampus, PCN = precuneus, ANG = angular gyrus.

#### Selective to the Posterior Medial Network?

To assess the selectivity of enhanced functional coupling within the posterior medial network for the Scrambled-Fixed clip, we conducted two additional analyses. First, we examined intra-subject functional correlation between the precuneus and a control region in early visual cortex (Figure 6). Unlike the posterior medial network, there were no changes in intra-subject functional correlation between the precuneus and early visual cortex over repetitions in any condition (all *ps* > .46). Moreover, there were no main effects of repetition or condition, nor repetition by condition interactions in either ANOVA (i.e., repetitions 1 vs. 6 and repetitions 1 through 6, all *ps* > .15).

**Figure 6.**
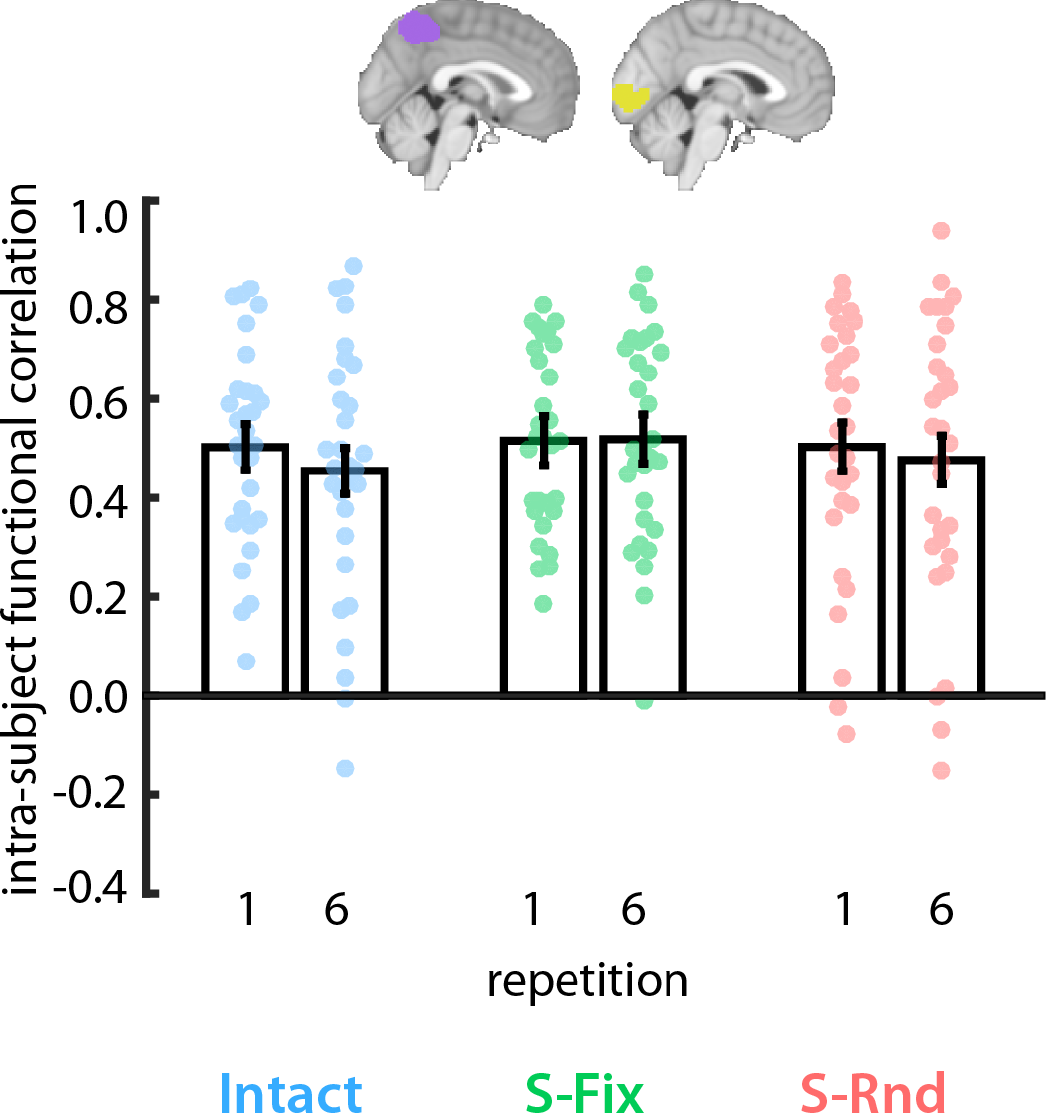
Functional correlation between the precuneus and early visual cortex. Intra-subject functional coupling between precuneus and visual cortex did not increase over repetitions in any condition. Dots are individual subjects. Error bars are ± 1 SEM of the within-subject difference between repetitions.

Next, we conducted whole-brain analyses of intra-subject functional correlation with the precuneus as a seed ROI, for the first and last repetitions of the Intact, Scrambled-Fixed, and Scrambled-Random clips. There were no reliable increases in intra-subject functional correlation for repetition 1 vs. 6 of the Intact or Scrambled-Random clips, correcting for multiple comparisons. In contrast, for the Scrambled-Fixed clip, significant increases in intra-subject functional correlation was observed in the hippocampus, angular gyrus, and medial parietal cortex (Figure 7), consistent with the ROI analyses of the posterior medial network. Although the medial parietal cluster overlaps with the seed ROI, this result indicates *changes* in intra-subject functional correlation over repetitions and thus is not guaranteed.

**Figure 7.**
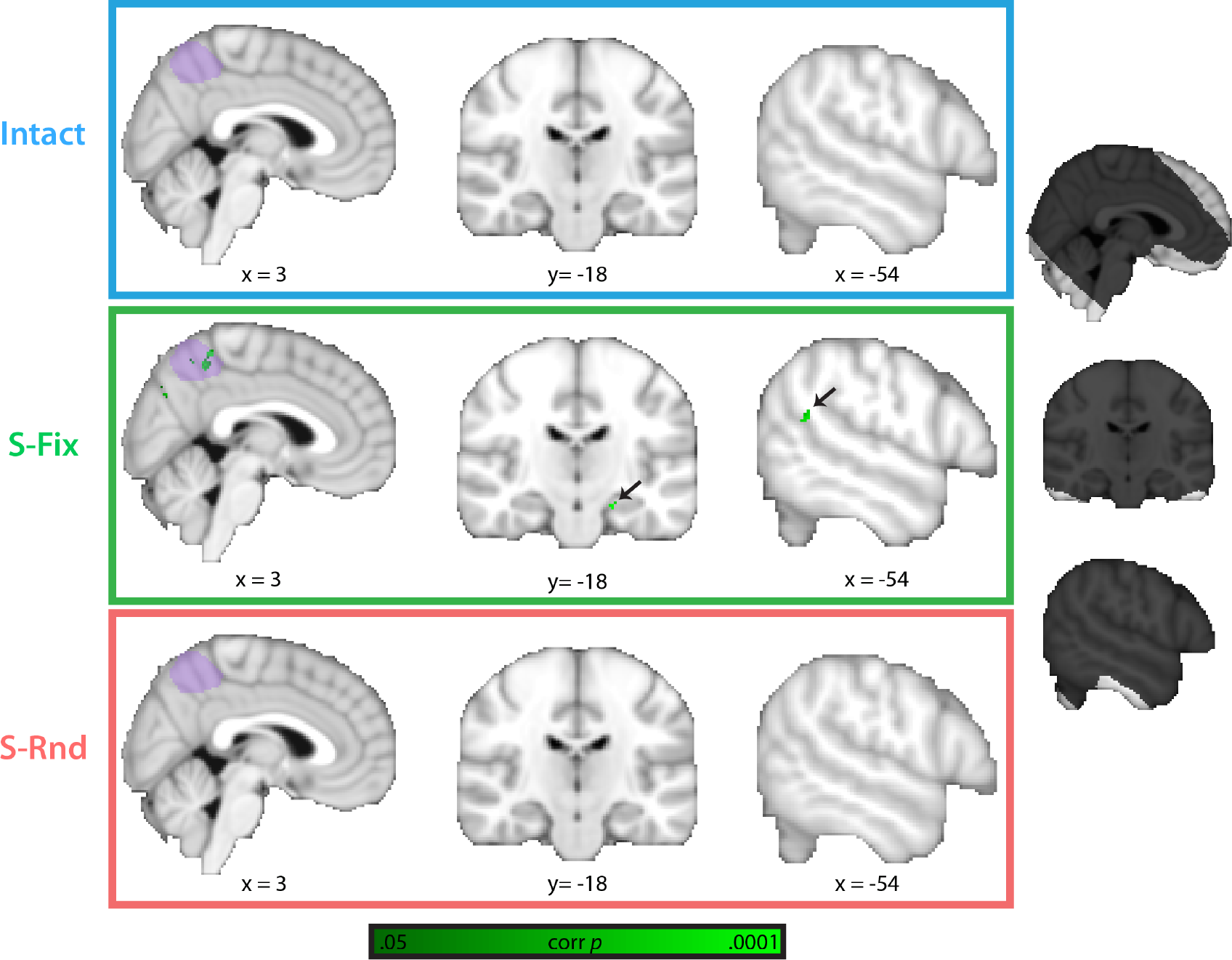
Changes in whole-brain precuneus (intra-subject) functional correlations from first to last repetition. Contrast depicts regions that increase in functional coupling with the precuneus (purple) from the first to last repetition of each clip (p < .05 corrected). No regions showed enhanced intra-subject functional correlation for the Intact (top) and Scrambled-Random (bottom) clips. For the Scrambled-Fixed clip (middle), greater precuneus intra-subject functional correlation (in green) was observed in the hippocampus and angular gyrus (and a medial parietal region that overlapped with the seed). The black mask (right) indicates to which voxels this analysis was applied (where > 90% of subjects had coverage).

#### What Does Enhanced Functional Coupling Reflect?

What is being learned in the Scrambled-Fixed condition that increases functional coupling in the posterior medial network? One possibility is that stronger intra-subject functional correlation reflects the regions’ activity becoming more locked to the stimulus. That is, upon initial exposure, the Scrambled-Fixed clip may seem incomprehensible and unpredictable, and thus is not consistently processed by different regions within the network. The extraction of temporal regularities over repetitions may allow posterior medial regions to focus on the same, regular aspects of the stimulus. This explanation can be tested by examining whether components of functional correlations that are locked to the stimulus increase over repetitions of the movie clips. Such stimulus-locked components can be measured with inter-subject functional correlation (Simony et al., 2016). In inter-subject functional correlation analyses (Figure 2D), correlations are calculated between the timecourse of activity in a given region in one subject and the timecourse of activity in a different region, averaged over all other subjects. As with inter-subject correlations (the same region between subjects), inter-subject functional correlation analysis isolates dynamics of brain activity attributable to the external stimulus viewed by all subjects, as opposed to components that are idiosyncratic to each individual or asynchronous across individuals. This is because aspects of between-region functional coupling that are related to the idiosyncratic thoughts of a particular individual will be removed when comparing the timecourse of activity between brain regions in *different* individuals. The commonality between different participants is that they are watching the same movie unfold over time. Thus, synchronized activity between different brain regions in different participants should be related to the common movie content that is being viewed, as opposed to individual-specific thoughts. In that sense, brain activity is locked to the stimulus because it is unfolding in the same way over time in different individuals watching the same movie.

We calculated inter-subject functional correlation between all pairs of posterior medial network ROIs and found no increase across repetitions in any condition (Supplementary Figure S4). Indeed, the only robust effects were *decreases* from repetition 1 to repetition 6 for HPC-PCN (Intact: *r*_Rep1_ = 0.31, *r*_Rep6_ = 0.07, *p* =0 from permutation test, 95% bootstrapped CI: 0.19–0.30; Scrambled-Fixed: *r*_Rep1_ = 0.11, *r*_Rep6_ = 0.03, *p* = .04 from permutation test, 95% bootstrapped CI: 0.007–0.16), PCC-PCN (Intact: *r*_Rep1_ = 0.37, *r*_Rep6_ = 0.12, *p* = 0 from permutation test, 95% bootstrapped CI: 0.16–0.36), and HPC-PCC (Intact: *r*_Rep1_ = 0.21, *r*_Rep6_ = 0.03, *p* = 0 from permutation test, 95% bootstrapped CI: 0.12–0.28; Scrambled-Random: *r*_Rep1_ = 0.05, *r*_Rep6_ = −0.09, *p* = .002 from permutation test, 95% bootstrapped CI: 0.08–0.21). This pattern of results — increased network coupling for the Scrambled-Fixed condition within but not between subjects — indicates that learning in the posterior medial network did not increase how much its responses were locked to the external stimulus. Moreover, this lack of an increase for inter-subject between-region correlations mirrors the lack of an increase in inter-subject within-region correlations (Supplementary Figure S2).

An alternative explanation, consistent with the lack of inter-subject enhancements in correlated temporal dynamics, is that what is being learned may be idiosyncratic to individuals. For example, over repetitions, each individual may converge on a unique meaning of the Scrambled-Fixed clip or learn particular segment transitions. This account predicts that the temporal dynamics within a region will be more similar for clip presentations toward the end vs. beginning of the six repetitions. That is, many learning-related changes may occur early during exposure, with a particular mode of processing or interpretation of the movie arrived at by the last repetition of each movie clip. Thus, for the Scrambled-Fixed clip, the correlation between within-region temporal dynamics for repetitions 1 vs. 2 may be lower than that for repetitions 5 vs. 6. However, this was not found: the within-region intra-subject correlation was not different for repetitions 1 and 2 compared to repetitions 5 and 6 of the Scrambled-Fixed clip (ANG: *r*_Rep1-2_ = −0.02, *r*_Rep5-6_ = −0.04, *t*_29_ = 0.25, *p* = .80, 95% CI: −0.14–0.18; HPC: *r*_Rep1-2_ = 0.0003, *r*_Rep5-6_ = 0.02, *t*_29_ = 0.32, *p* = .75, 95% CI: −0.11–0.08; PCC: *r*_Rep1-2_ = 0.06, *r*_Rep5-6_ = −0.01, *t*_29_ = 1.05, *p* = .30, 95% CI: −0.08–0.24; PCN: *r*_Rep1-2_ = 0.15, *r*_Rep5-6_ = 0.08, *t*_29_ = 0.91, *p* = .37, 95% CI: −0.10–0.25).

There were also no differences between these repetition pairs for the Scrambled-Random clip (all *p*s > .62); no differences are expected here because the Scrambled-Random clip was viewed in a different order for each repetition, thus brain activity is being compared for objectively different stimuli.

For the Intact clip, we found greater similarity early on (i.e., the first two vs the last two presentations of the clip) in precuneus and hippocampus (precuneus: *r*_Rep1-2_ = 0.44, *r*_Rep5-6_ = 0.13, *t*_29_ = 5.33, *p* = .0001, 95% CI: 0.22–0.48; hippocampus: *r*_Rep1-2_ = 0.18, *r*_Rep5-6_ = 0.08, *t*_29_ = 2.34, *p* = .03, 95% CI: 0.13– 0.20). This effect for the Intact clip was not reliable in angular gyrus (*r*_Rep1-2_ = 0.19, *r*_Rep5-6_ = 0.08, *t*_29_ = 1.78, *p* = .09, 95% CI: −0.02–0.25) or posterior cingulate (*r*_Rep1-2_ = 0.17, *r*_Rep5-6_ = 0.08, *t*_29_ = 1.54, *p* = .13, 95% CI: - 0.03–0.24). These results are inconsistent with the idea of gradually arriving at a particular manner of processing the movie clip.

## Discussion

The brain contains a hierarchy of regions that respond to information over varying timescales (Chaudhuri et al., 2015; Hasson et al., 2008; Murray et al., 2014). Unlike sensory regions, in which activity at any given moment is predominantly driven by the immediate environment, higher-order regions in the posterior medial network integrate events over many seconds or minutes (Baldassano et al., 2017; Chen et al., 2016; Hasson et al., 2015; Simony et al., 2016). Prior studies have often explored temporal integration by analyzing within-region dynamics for naturalistic stimuli. The reliability of a region’s temporal dynamics is calculated either by examining how highly that region’s activity is correlated between different subjects watching the same stimulus, or within a subject over repetitions of the same stimulus (e.g., Hasson et al., 2004, 2008; Honey et al., 2012a,b; Lerner et al., 2011). These studies find that the reliability of temporal fluctuations in posterior medial regions, whether measured between subjects or within a subject, is highest for intact movies and stories and is reduced when these stimuli are temporally scrambled.

Higher within-region reliability for intact vs. scrambled stimuli may occur because intact movies and stories contain rich temporal structure that can exploit pre-existing event schemas or scripts (Barlett, 1932; Bower et al., 1979; van Kesteren et al., 2012): Over a lifetime, we have learned, via statistical learning, which events tend to follow each other when, for example, we walk into a lobby and push a button for an elevator. However, statistical learning can also take place much more rapidly, allowing temporal structure to be learned within a relatively short period of time (Aslin & Newport, 2012; Schapiro et al., 2017). Events with novel structure may initially violate expectations because they do not align with pre-existing schemas, but can become predictable after only a handful of repetitions (Turk-Browne et al., 2010). The goal of this study was to explore whether temporal structure is sufficient to engage the posterior medial network, even when it is not schema-consistent.

Learning of temporal structure by regions in the posterior medial network may not manifest as increases in the consistency of within-region dynamics across stimulus repetitions, however, because a region may become *less* similar to itself over stimulus repetitions if learning occurs in that region. Learning may instead be revealed as changes in between-region functional coupling: For example, the brain’s network dynamics change from the first to the second viewing of a movie (Andric et al., 2016). One possibility is that regions within a network may communicate more, or differently, with one another when a stimulus is recognized as potentially significant because it contains temporal structure. We therefore examined both within- and between-region dynamics to look for evidence of learning in the brain.

We found that temporal dynamics within posterior medial regions were more reliable for movie clips that were consistent vs. inconsistent with pre-existing schemas (also see Keidel et al., 2017). This was true both within individuals over clip repetitions and between individuals within repetition, replicating and extending prior work with intact vs. scrambled stimuli (e.g., Hasson et al., 2008; Honey et al., 2012a; Lerner et al., 2011). However, over repetitions of a scrambled movie with fixed temporal structure, intra-subject functional coupling (i.e., functional connectivity) between posterior medial regions gradually increased, rising to the level of the intact movie. Such increased coupling may be a signature of learning temporal structure in the brain, and might have been missed by prior studies because scrambled stimuli were presented at most twice in the same order, and functional correlations in the posterior medial network were not examined (e.g., Hasson et al., 2008; Honey et al., 2012a). Thus, examining within-region dynamics may not be sufficient to reveal learning: Between-region functional coupling can show increases in network engagement when within-region dynamics suggest no such effect.

### What Is Enhanced Network Coupling Reflecting?

Although enhancements in posterior medial network intra-subject functional correlation was strongest for the Scrambled-Fixed clip, a subset of region pairs showed enhanced coupling over repetitions for the Intact and/or Scrambled-Random clips. Such increases may occur because there is some temporal structure to be learned even here. In the Intact clip, information about the specific sequence of events can be learned when the movie is watched several times. That is, while individuals can draw upon event schemas to make sense of what is happening in a general sense for the intact movie, repeated viewings allow learning of movie-specific information, e.g., which characters interact in what order, and how they move over space and time. Indeed, subjects’ memory recalls indicated that they did learn the sequence of events. In the Scrambled-Random clip, there was also the possibility of extracting some structure over repetitions, as regularities existed on a short timescale *within* each segment that comprised the clip. Nevertheless, the Scrambled-Fixed clip offered the most opportunity for learning: In addition to what could be learned in the Scrambled-Random clip, there were deterministic transitions between clip segments, which were unknown on the first repetition. Long time-scale temporal structure may therefore be key for engaging the posterior medial network, even if it is inconsistent with pre-existing event schemas.

Another possibility is that the enhanced functional coupling is reflecting greater comprehension of the narrative of each movie clip, instead of learning of temporal structure. Indeed, all three clips were understood by subjects, as evidenced in their memory recall at the end of the experiment. After six repetitions, subjects were able to recall many details about each movie, with memory slightly better for the Intact vs Scrambled-Fixed and Scrambled-Random clips, and memory for the Scrambled-Fixed and Scrambled-Random clips not differing (see Behavioral Results). Moreover, subjects tended to structure their recall in the order of the intact narrative, even for the Scrambled-Fixed and Scrambled-Random clips (i.e., they were able to reorganize the scrambled segments into a coherent narrative).

We think this is a possible contributing factor to enhanced functional coupling, and may account for why there were some increases in functional coupling for the Scrambled-Random clip. However, we think it is unlikely that this is the full explanation, as enhancements in functional coupling were numerically strongest and most consistent for the Scrambled-Fixed clip (Figures 4, 5, 7). Thus, although there may be a role for narrative comprehension, our data suggest that there is an additional role for temporal structure.

To gain more insight into this question, future work could explore changes in functional coupling while collecting online behavioral measures of narrative comprehension and temporal structure learning (i.e., behavioral tests between each clip repetition). In that way, the changes in functional coupling can be linked to changes in these behavioral measures over stimulus repetitions. Our behavioral data were collected at the end of all six repetitions and we did not directly test learning of temporal structure; thus, the current study cannot adjudicate between these possibilities.

What is the content of functionally coupled representations in the posterior medial network? That is, as intra-subject functional correlation increases over the course of exposure to temporally structured but scrambled clips, what do posterior medial regions come to jointly represent? We examined two possibilities — increased stimulus locking and converging idiosyncratic interpretations — but found no evidence for either. That is, posterior medial network dynamics were largely unique to each individual and repetition rather than shared across individuals or repetitions. For example, inter-subject correlation and inter-subject functional correlation, used to isolate stimulus-locked temporal fluctuations, either decreased or were unaffected (Supplementary Figure S2 and S4). Moreover, within-region, intra-subject temporal dynamics were no more similar for clip presentations at the end vs. beginning of clip repetitions, suggesting that individuals did not gradually arrive at a particular mode of processing the movie clips. Thus, network activity was not strictly tied to the stimulus that was commonly viewed by subjects, but was related in part to idiosyncratic cognitive processes for each individual and even for each repetition for a given individual.

What is enhanced functional coupling reflecting then? One possibility is that the network becomes more engaged, but in different ways over repetitions, when a stimulus has been recognized as meaningful, perhaps as a result of detecting temporal structure from prior experience. For example, what is attended may change on each repetition of the clip as a result of prior statistical learning, helping to enable the extraction of new information (Zhao et al., 2013). These switches in attention may manifest as enhanced functional coupling between regions that is dissociable from stimulus-locked correlations. Indeed, such “background” correlations (i.e., task-based functional coupling not attributable to stimuli) have been observed in the visual system when attention switches between objects (Al-Aidroos et al., 2012; Córdova et al., 2016). A similar mechanism may operate in the posterior medial network, allowing us to learn novel, episode-specific details in a familiar event schema. This could explain how, for example, we can remember unique moments from the last time we went to a restaurant, as opposed to only the moments common to all restaurant visits.

A study from van Kesteren and colleagues (2010) offers further insight about what enhanced functional correlations for temporally structured events could mean. During viewing of an intact movie’s conclusion, coupling between the hippocampus and ventromedial prefrontal cortex was higher in individuals who had viewed a scrambled version of the preceding part of the movie, relative to individuals who had viewed the intact version of the preceding part (which provided schematic structure). The authors suggest that functional coupling between hippocampus and ventromedial prefrontal cortex might be enhanced when there is greater difficulty integrating novel information because pre-existing schema are not available. Intriguingly, this was observed alongside higher correlations in inter-subject temporal dynamics in the ventromedial prefrontal cortex for the schema consistent vs. inconsistent group.

We were not able to examine the ventromedial prefrontal cortex in the current study (because of signal dropout, see Figure 7), but this perspective would predict reductions in functional coupling between the hippocampus and ventromedial prefrontal cortex over repetitions of the Scrambled-Fixed clip, because with more exposure to temporal structure, the ease of integrating information may increase (i.e., a new schema potentially begins to form). Such a decrease in functional correlation would serve as a contrast to the increasing posterior medial network coupling found in the current study, and would provide a dissociation among different components of the default mode network (e.g., Andrews-Hanna et al., 2010).

An alternative perspective is that the decrease in inter-subject correlation over repetitions, both within-region (Supplementary Figure S2) and between regions (Supplementary Figure S4) indicates that subjects are gradually becoming disengaged from the movies being viewed. If so, increases in within-subject functional correlations might just indicate greater disengagement of the “default mode” (Raichle, 2015) from the external world, as subjects become fatigued and turn to internal thoughts rather than processing external stimulus. However, if this were the case, such increased functional correlation should occur for all three clips being viewed and for an equivalent amount, rather than being largest for the Scrambled-Fixed clip. Moreover, if high functional correlation between posterior medial regions indicates disengagement from external stimuli because of fatigue, then it is not clear why such high correlations were observed on the very first exposure to the Intact movie clip (Figure 4).

One might argue that subjects disengage immediately, on the very first viewing of the Intact clip, because moment-by-moment actions are relatively predictable (i.e., if a character is walking toward an elevator, he will likely enter the elevator and the door will likely close after him). However, if subjects disengaged from the start, one might expect their memory for the Intact clip to be worse than that for the scrambled clips, but that is not the case – memory was *better* for the Intact clip (see Behavioral Results). Moreover, if individuals were disengaging immediately for the Intact clip, then it is not clear why both intra-subject and inter-subject correlations (Figure 3) were higher for the Intact vs scrambled clips. That is, it is hard to explain synchronized activity between individuals (Figure 3B) if each individual was disengaged from the movie clips and instead turned to idiosyncratic internal thoughts. Likewise, one would have to argue that a given individual’s internal thoughts and their temporal trajectory were stable from the first to the last repetition of the Intact clip (Figure 3A).

That said, we cannot rule out differential engagement for different movie clips because we did not have online behavioral measures of attention. One way to measure online attention would be to have participants respond with a button press each time a particular character is seen or event occurs. Future studies can include such behavioral tasks to assess engagement online, and relate this to changes in posterior medial network coupling.

### Temporal Sequence Learning in the Hippocampus

The increased coupling between the hippocampus and other regions in the posterior medial network over repetitions of temporally structured, scrambled clips contributes to the literature on time and sequence coding in hippocampus (e.g., DuBrow and Davachi, 2014, 2016; Hsieh et al., 2014; MacDonald et al., 2011; Mankin et al., 2012; Manns et al., 2007; Pastalkova et al., 2008; see Davachi and DuBrow, 2015; Eichenbaum, 2013; Ranganath and Hsieh, 2015) and studies showing a role for the hippocampus in memory for intact movies (e.g., Chen et al., 2017; Lehn et al., 2009; Gelbard-Sagiv et al., 2008). This is also consistent with studies of statistical learning that have found learning-related changes in hippocampal activity (Bornstein & Daw, 2013; Hindy et al., 2016; Schapiro et al., 2012; Turk-Browne et al., 2009, 2010). The stimuli in these studies were groupings of shapes, objects, faces, and/or scenes, showing that the hippocampus can learn temporal regularities even among arbitrary stimuli that cannot exploit pre-existing schemas. Our results complement this work by showing changes in hippocampal coupling with other brain regions during repeated exposure to naturalistic temporal regularities.

The statistical learning literature emphasizes that a key consequence of extracting temporal structure is the ability to generate accurate predictions (e.g., Schapiro et al., 2012; Turk-Browne et al., 2010). Indeed, the hippocampus exhibits anticipatory signals that can impact goal-directed behavior (e.g., Bornstein and Daw, 2013; Brown et al., 2016; Hindy et al., 2016; Johnson and Redish, 2007; Pfeiffer and Foster, 2013; for review, see Buckner, 2010; Redish, 2016). Given the increased coupling of posterior medial regions with the hippocampus, one speculation is that this may be contributing to hippocampal prediction. For example, integration of long-timescale information in posterior medial regions may allow the hippocampus to generate predictions about events in the more distant future. Indeed, providing such a temporal history to the hippocampus helps it extract regularities and generate predictions (Schapiro et al., 2017). Another possibility is the flow of information goes the other way: that is, predictive signals in the hippocampus may be communicated to posterior medial regions to help them with temporal integration. Our data cannot adjudicate between these possibilities, as we cannot measure the directionality of information flow, but these remain interesting questions for future research with other methods (e.g., intracranial EEG).

### Concluding Remarks

In conclusion, the posterior medial network is critical for remembering event sequences and integrating information over long timescales (Hasson et al., 2015; Ranganath and Ritchey, 2012). We found that repeated viewing of novel temporal sequences leads to increased functional coupling between the precuneus, hippocampus, angular gyrus, and posterior cingulate cortex. This occurred in the absence of evidence for increased stimulus-locked responses, suggesting idiosyncratic and labile temporal dynamics. These findings highlight how repeated exposure to temporal structure can induce changes in network organization over time, which may be a necessary precursor for learning.

## Acknowledgments

This work was funded by NIH grants R01-EY021755 to N.B.T.B. and R01-MH112357-01 to U.H. This project was also made possible through the support of a grant from the John Templeton Foundation. The opinions expressed in this publication are those of the authors and do not necessarily reflect the views of these agencies. The authors have no competing interests to declare.

## Conflict of Interest

None

## Author Contributions

Mariam Aly: Conceptualization, Methodology, Validation, Formal analysis, Investigation, Data curation, Writing – original draft preparation, Writing – review & editing, Visualization, Project administration. Janice Chen: Conceptualization, Methodology, Investigation, Writing – review & editing, Project administration.

Nicholas Turk-Browne: Conceptualization, Methodology, Resources, Writing – review & editing, Supervision, Funding acquisition

Uri Hasson: Conceptualization, Methodology, Resources, Writing – review & editing, Supervision, Funding acquisition

## Supplementary Information

**Figure S1.**
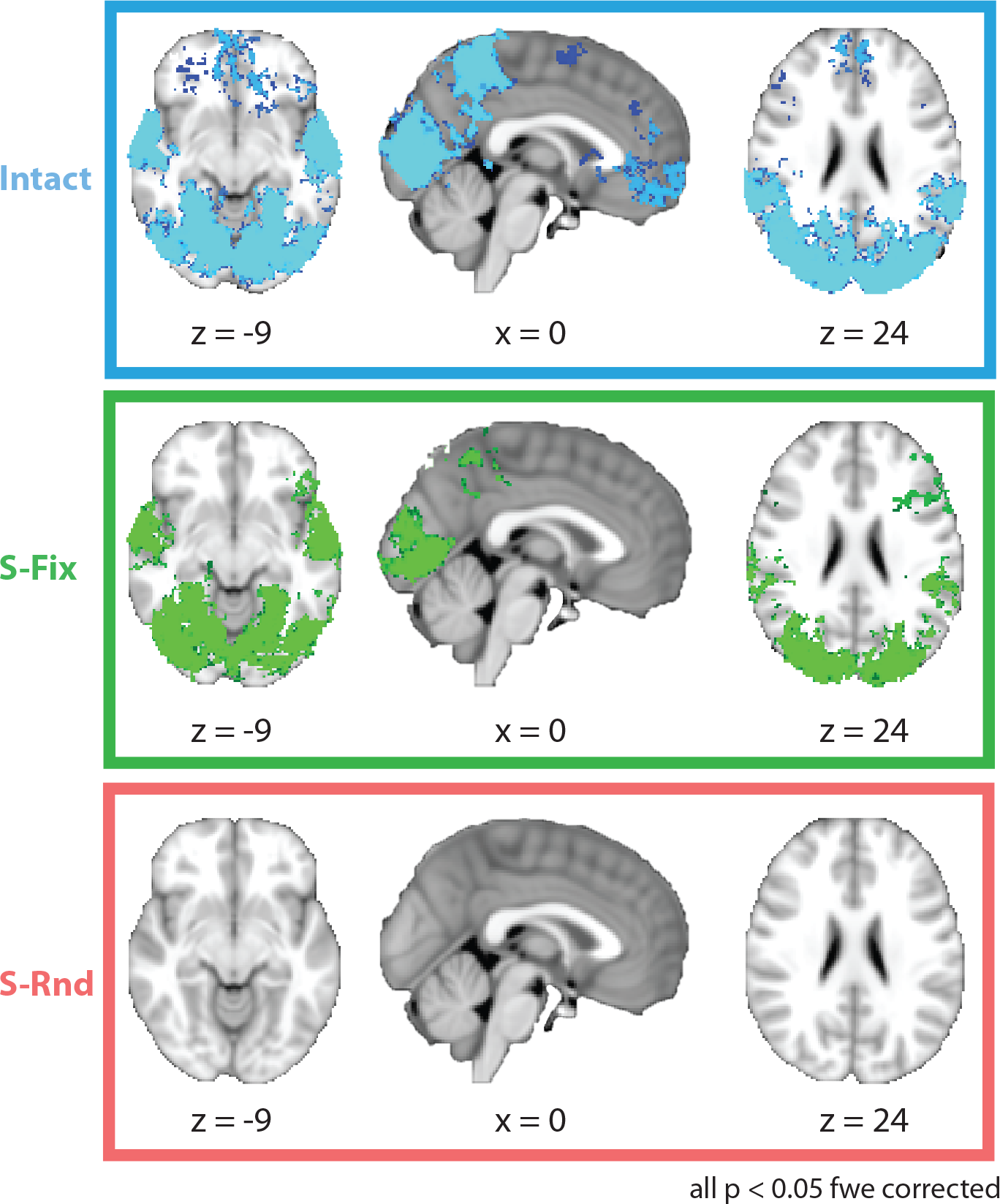
Whole-brain analysis of intra-subject temporal dynamics for clip repetition 1 vs clip repetition 6. We calculated, for each voxel in the brain for each individual, the correlation between the timecourse of activity for repetition 1 of each clip and repetition 6 for each clip. These whole-brain intra-subject correlation maps were analyzed at the group level using random-effects non-parametric tests (randomise in FSL), and corrected for family-wise error (FWE) across all voxels. A number of regions in occipitotemporal cortex, parietal cortex, and prefrontal cortex showed statistically significant intra-subject correlations between repetition 1 and repetition 6 of the Intact and Scrambled-Fixed clips. This was not the case for the Scrambled-Random clip, in which no statistically significant results were obtained. This was expected, because the stimuli differed from repetition to repetition for the Scrambled-Random, but not the Intact or Scrambled-Fixed, movie.

**Figure S2.**
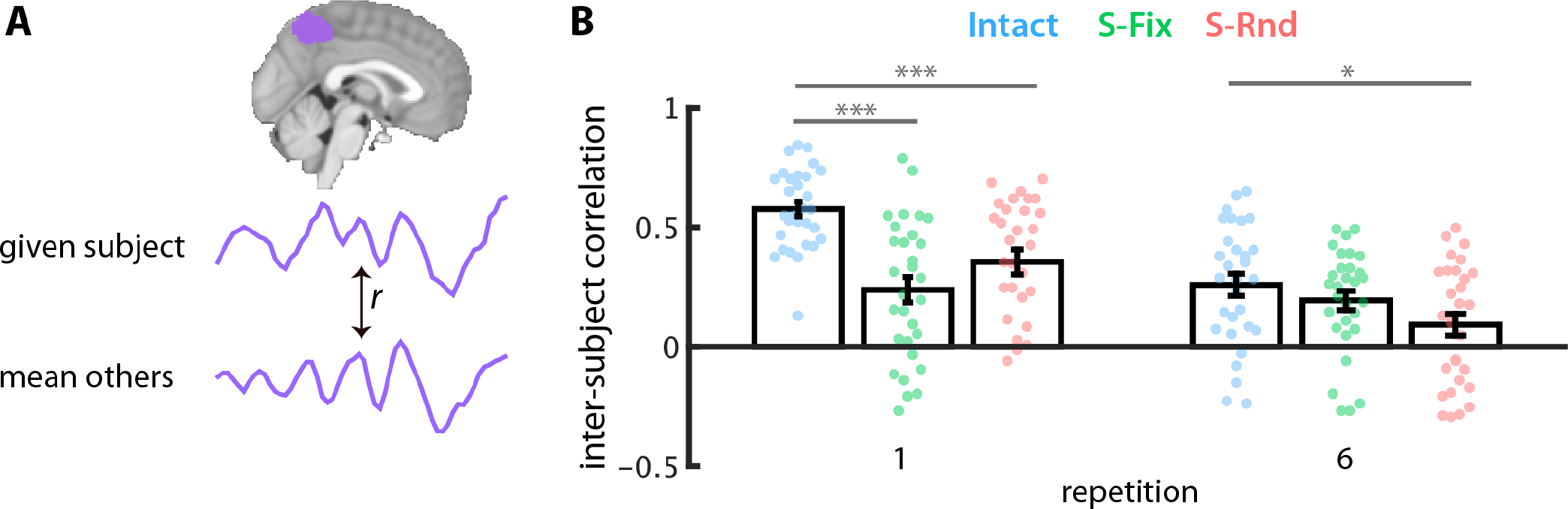
Inter-subject temporal dynamics across clip repetitions. **(A)** Response reliability was assessed by correlating, for the first and last repetition separately, each subject’s timecourse of BOLD activity in the precuneus with the mean precuneus timecourse across all other subjects (i.e., inter-subject correlation). **(B)** Statistical analyses for repetition 1 are reported in the main text; the data are shown here again for comparison to repetition 6. Inter-subject correlation was higher for repetition 1 vs. repetition 6 of the Intact clip (*r*_Rep1_ = 0.58, *r*_Rep6_ = 0.26; 95% bootstrapped CI: 0.30–0.51; *p* = 0 from permutation test). Inter-subject correlation was also higher for repetition 1 vs. repetition 6 of the Scrambled-Random clip (*r*_Rep1_ = 0.36, *r*_Rep6_ = 0.09; 95% bootstrapped CI: 0.13–0.45; *p* = .002 from permutation test). In contrast, repetition 1 and repetition 6 were not different from one another for the Scrambled-Fixed clip (*r*_Rep1_ = 0.24, *r*_Rep6_ = 0.19; 95% bootstrapped CI: −0.07–0.21; *p* = .36 from permutation test). For repetition 6, inter-subject correlation was reliably above zero for all three conditions (Intact: *r*_Rep6_ = 0.26; 95% bootstrapped CI: 0.18–0.38; *p* = 0 from permutation test; Scrambled-Fixed: *r*_Rep6_ = 0.19; 95% bootstrapped CI: 0.12– 0.28; *p* = 0 from permutation test; Scrambled-Random: *r*_Rep6_ = 0.09; 95% bootstrapped CI: 0.006–0.18; *p* = .04 from permutation test). Correlations were higher for the Intact clip vs. the Scrambled-Random clip on repetition 6 (95% bootstrapped CI: 0.04–0.32; *p* = .02 from permutation test). The Intact and Scrambled-Fixed clips were not different from one another on repetition 6 (95% bootstrapped CI: −0.04–0.20; *p* = .22 from permutation test), and neither were the Scrambled-Fixed and Scrambled-Random clips (95% bootstrapped CI: −0.008–0.21; *p* = .12 from permutation test). Dots indicate individual subjects. Error bars are ± 1 SEM. * *p* < .05, ** *p* < .01, *** *p* < .001.

**Figure S3.**
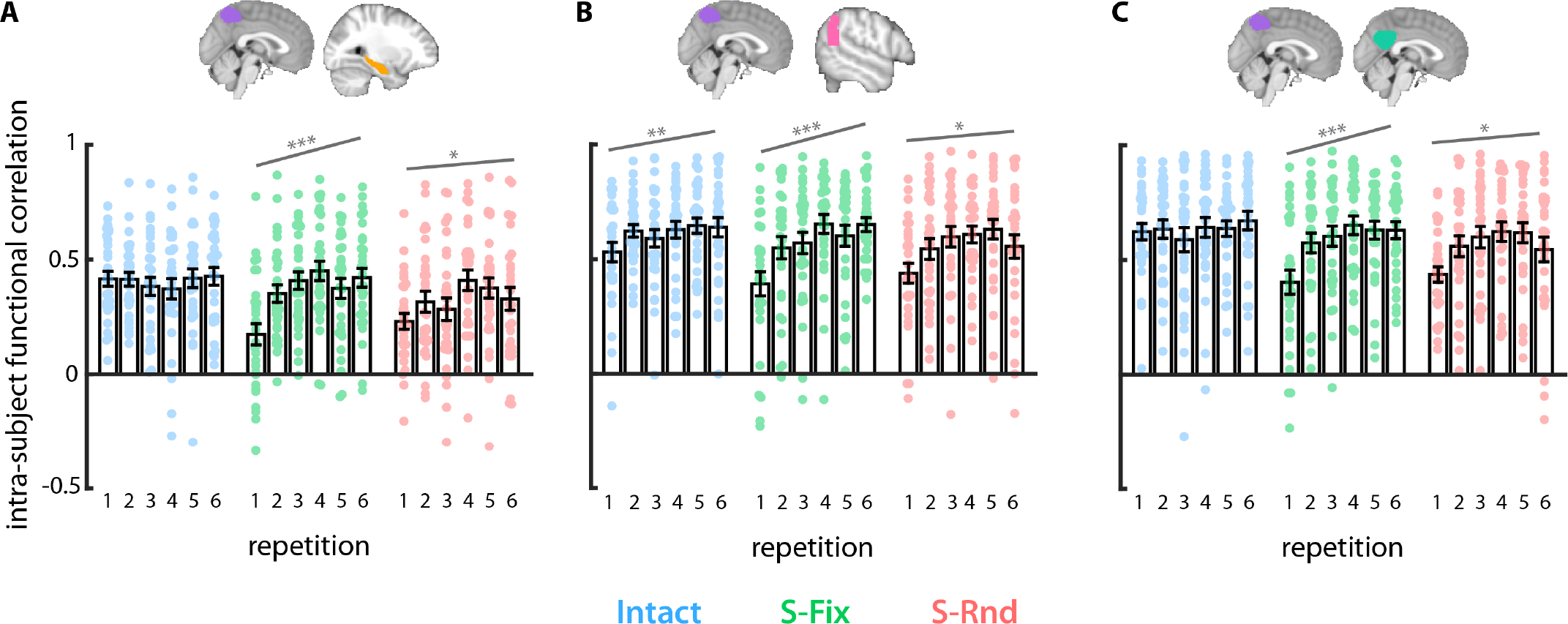
Intra-subject functional correlation across all six repetitions of each clip. **(A)** Intra-subject functional correlation between precuneus and hippocampus increased monotonically across all six repetitions of the Scrambled-Fixed clip. (β = 0.05, *t*_29_ = 4.06, *p* = .0003, 95% CI: 0.02–0.07). The slope was not different from zero for the Intact clip (β = 0.004, *t*_29_ = 0.43, *p* = .67, 95% CI: −0.02–0.02) and was numerically smaller for the Scrambled-Random clip (β = 0.03, *t*_29_ = 2.17, *p* = .04, 95% CI: 0.002–0.06). **(B)** Intra-subject functional correlation between precuneus and angular gyrus increased monotonically across all six repetitions of the Scrambled-Fixed clip (β = 0.07, *t*_29_ = 4.68, *p* = .00006, 95% CI: 0.04–0.09). There were also numerically smaller increases for the Intact (β = 0.04, *t*_29_ = 3.43, *p* = .002, 95% CI: 0.02–0.06) and Scrambled-Random clips (β = 0.04, *t*_29_ = 2.42, *p* = .02, 95% CI: 0.007–0.08). **(C)** Intra-subject functional correlation between precuneus and posterior cingulate cortex increased monotonically for the Scrambled-Fixed clip (β = 0.06, *t*_29_ = 4.76, *p* = .00005, 95% CI: 0.03–0.08). There was a numerically smaller increase for the Scrambled-Random clip (β = 0.04, *t*_29_ = 2.48, *p* = .02, 95% CI: 0.007–0.08). The slope for the Intact clip was not different from zero (β = 0.02, *t*_29_ = 1.58, *p* = .12, 95% CI: −0.006–0.05). We also fit quadratic and logarithmic functions to the data, but the results were generally consistent with the linear fits (i.e., there was only one case where the data were a statistically significant fit to one function but not the others: coupling between the angular gyrus and the precuneus increased over repetitions for the Intact clip when fit to linear and logarithmic functions, but not a quadratic function). Dots indicate individual subjects. Error bars are ± 1 SEM. * *p* < .05, ** *p* < .01, *** *p* < .001.

**Figure S4.**
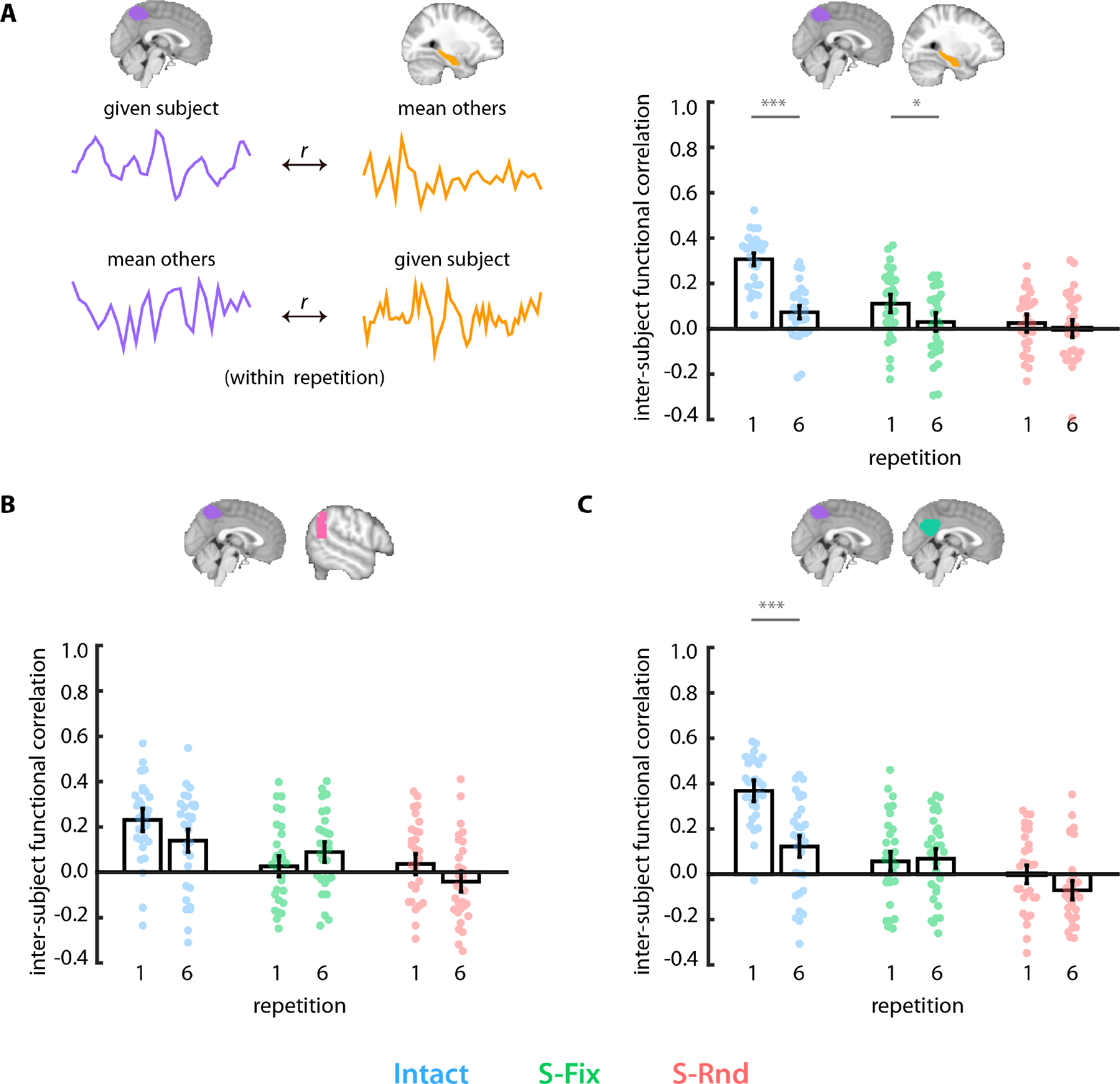
Inter-subject functional correlation. **(A)** The timecourse of BOLD activity in the precuneus for a given subject was correlated with the mean timecourse of BOLD activity in the hippocampus for all other subjects. The timecourse of BOLD activity in the hippocampus for that same subject was also correlated with the mean timecourse of BOLD activity in the precuneus for all subjects. The average of these correlations was the inter-subject functional correlation value for the hippocampus and the precuneus for that subject. Inter-subject functional correlation decreased from repetition 1 to repetition 6 of the Intact clip (*r*_Rep1_ = 0.31, *r*_Rep6_ = 0.07, *p* =0 from permutation test, 95% bootstrapped CI: 0.19–0.30) and the Scrambled-Fixed clip (*r*_Rep1_ = 0.11, *r*_Rep6_ = 0.03, *p* = .04 from permutation test, 95% bootstrapped CI: 0.007–0.16), and did not change for the Scrambled-Random clip (*r*_Rep1_ = 0.03, *r*_Rep6_ = 0.001, *p* = .49 from permutation test, 95% bootstrapped CI: −0.06–0.11). **(B)** Inter-subject functional correlation between the precuneus and angular gyrus did not change over repetitions in any condition (Intact: *r*_Rep1_ = 0.23, *r*_Rep6_ = 0.14, *p* = .09 from permutation test, 95% bootstrapped CI: −0.006–0.20; Scrambled-Fixed: *r*_Rep1_ = 0.03, *r*_Rep6_ = 0.09, *p* = .16 from permutation test, 95% bootstrapped CI: −0.15–0.02; Scrambled-Random: *r*_Rep1_ = 0.04, *r*_Rep6_ = −0.04, *p* = .15 from permutation test, 95% bootstrapped CI: −0.008–0.17). **(C)** Inter-subject functional correlation between the precuneus and posterior cingulate cortex decreased over repetitions for the Intact clip (*r*_Rep1_ = 0.37, *r*_Rep6_ = 0.12, *p* = 0 from permutation test, 95% bootstrapped CI: 0.16–0.36), but did not change over repetitions of the Scrambled-Fixed clip (*r*_Rep1_ = 0.06, *r*_Rep6_ = 0.07, *p* = .77 from permutation test, 95% bootstrapped CI: −0.09–0.07) or the Scrambled-Random clip (*r*_Rep1_ = −0.0005, *r*_Rep6_ = −0.07, *p* = .10 from permutation test, 95% bootstrapped CI: −0.01–0.14).

### Supplementary Methods

Below are brief descriptions of the three movie clips used from *The Grand Budapest Hotel*.

Clip A was an interview scene between a hotel concierge and a boy applying for the position of lobby boy. The prospective lobby boy is asked a series of questions about his experience, education, family, and desire for the job as he and the concierge move through the long lobby of the Grand Budapest Hotel. The concierge is repeatedly interrupted by hotel staff and guests as he walks through the lobby. These individuals ask various questions or receive comments from the concierge. The clip concludes with the prospective lobby boy and concierge walking into an elevator, going to another floor, and walking into a room in which the concierge picks up an envelope from a table.

Clip B was a scene depicting a theft of a painting by the hotel concierge and the lobby boy (the boy who was interviewed in clip A). The hotel concierge and the lobby boy are discussing a painting while seated in front of a window, through which a kitchen and kitchen employees can be seen. The hotel concierge is describing the painting to the lobby boy, who interrupts to ask if he can see the painting. The hotel concierge acquiesces, and the two of them walk through the hotel, up some stairs, and into a large room in which the painting is prominently displayed. They look at the painting for a while, before silently agreeing to steal it. The lobby boy brings a stool to the concierge, who stands on the stool and lifts the painting from the wall.

Clip C was a chase scene between a bakery girl who had obtained the painting (that was stolen in Clip B) and a man who believed he was the rightful owner of said painting. The man saunters into the hotel with his sisters and is greeted at the door. While being greeted in the lobby and provided with information about his room, he notices the bakery girl on the hotel stairs, with a package under her arm, which he took to be his painting. The bakery girl notices him and starts to run away, up the stairs and to an elevator. The man catches up to her in the elevator, and the scene ends as they are eyeing each other. Interleaved with this narrative, the lobby boy and concierge arrive at the hotel dressed as bakers, bribe a man at the front desk with baked goods, and rush through the hotel looking for the bakery girl.

## Footnotes

1 “Learning” can refer to changes in behavior as a result of experience, or changes in the brain as a result of experience – with these latter changes ultimately having effects on cognition and behavior. Because the current study did not include behavioral measures of learning temporal structure, our assessments of learning are based only on changes in brain activity with experience.

2 “Repetition 1” here and elsewhere refers to the first time a movie clip was viewed. We clarify this because “Repetition 1” may be otherwise interpreted as the *second* time a clip was viewed.

